# Replicability of spatial gene expression atlas data from the adult mouse brain

**DOI:** 10.1101/2020.10.16.343210

**Authors:** Shaina Lu, Cantin Ortiz, Daniel Fürth, Stephan Fischer, Konstantinos Meletis, Anthony Zador, Jesse Gillis

## Abstract

**Background:** Spatial gene expression is particularly interesting in the mammalian brain, with the potential to serve as a link between many data types. However, as with any type of expression data, cross-dataset benchmarking of spatial data is a crucial first step. Here, we assess the replicability, with reference to canonical brain sub-divisions, between the Allen Institute’s *in situ hybridization* data from the adult mouse brain (ABA) and a similar dataset collected using Spatial Transcriptomics (ST). With the advent of tractable spatial techniques, for the first time we are able to benchmark the Allen Institute’s whole-brain, whole-transcriptome spatial expression dataset with a second independent dataset that similarly spans the whole brain and transcriptome.

**Results:** We use LASSO, linear regression, and correlation-based feature selection in a supervised learning framework to classify expression samples relative to their assayed location. We show that Allen reference atlas labels are classifiable using transcription, but that performance is higher in the ABA than ST. Further, models trained in one dataset and tested in the opposite dataset do not reproduce classification performance bi-directionally. Finally, while an identifying expression profile can be found for a given brain area, it does not generalize to the opposite dataset.

**Conclusions:** In general, we found that canonical brain area labels are classifiable in gene expression space within dataset and that our observed performance is not merely reflecting physical distance in the brain. However, we also show that cross-platform classification is not robust. Emerging spatial datasets from the mouse brain will allow further characterization of cross-dataset replicability.

## Background

Over a decade ago, the first whole-transcriptome, spatially-resolved gene expression dataset from the adult mouse brain was collected by the Allen Institute using *in situ* hybridization (ABA) (1,2). Since its release, this dataset has become a cornerstone for modern neurobiologists who often use it as a first point of reference for gene expression in the mouse brain. The generation of this dataset was a laborious effort requiring many years, the work of many scientists, and many sacrificed mice. Recently, however, new techniques have made it possible to easily sequence whole transcriptomes while retaining fine-scale spatial information (3–6). These new techniques offer a more high-throughput and finer spatial resolution than sequencing multiple areas using traditional RNA sequencing following microdissection. This influx of technologies preserving spatial origin of transcripts presents the opportunity to benchmark the ABA data for the first time.

Validating spatially-located expression data is especially important in neuroscience where it has the potential to serve as a link between the molecular, meso-scale, and emergent properties of the brain such as gene expression, circuitry, and behavior, respectively (1). Emerging experimental approaches (7–9) and techniques (10–14) have already begun to link multi-source information from the mouse brain. In order to perform robust multi-modality studies, we must first benchmark within one type of data. As the sole reference spatial dataset, benchmarking the ABA data is essential to assess the robustness of the observed gene expression patterns across distinct experiments and technological platforms. Obtaining replicable results across gene expression assays is notoriously challenging, so cross-platform, cross-dataset transcriptomics benchmarking has proved crucial since early transcriptome assays in the form of microarrays (15,16).

To address this need for spatial transcriptomics and cross-modality robustness in the brain, here we undertook a whole-brain benchmarking of the ABA via linking gene expression and anatomy. We analyzed a spatial gene expression dataset from one adult mouse brain collected using spatial transcriptomics (ST) (17) (see methods) alongside the ABA. To assess replicability of spatial gene expression with these two datasets, we could look:

1. within brain, within technique to assay the associated noise level of a technique;
2. within brain, across technique to assay the relative biases of each technique;
3. across brain, within technique to assay batch effects of tissue processing, alignment across the brain, and biological variability;
4. across brain, across technique which would have noise from 2 and 3; or
5. across brain, across technique with reference to named brain areas to assay noise as in 4 in addition to variability from brain area segmentation and naming.

Focusing on biological conclusions that could be drawn from replicable spatial data, we turn our attention to variability across brain and across technique relative to brain areas, as described in 5. We principally ask if canonical, anatomically-defined brain areas from the Allen Reference Atlas (ARA) can be identified using gene expression alone and, in corollary, how well this replicates across the ABA and ST datasets. We use an interpretable supervised learning framework for classification, where the target values are the ARA brain area labels and the features are the gene expression profiles for samples from across the whole brain (Figure 1a, b). We choose to use linear modeling to maintain easily interpretable models that can be related to underlying biology.

**Figure 1.**
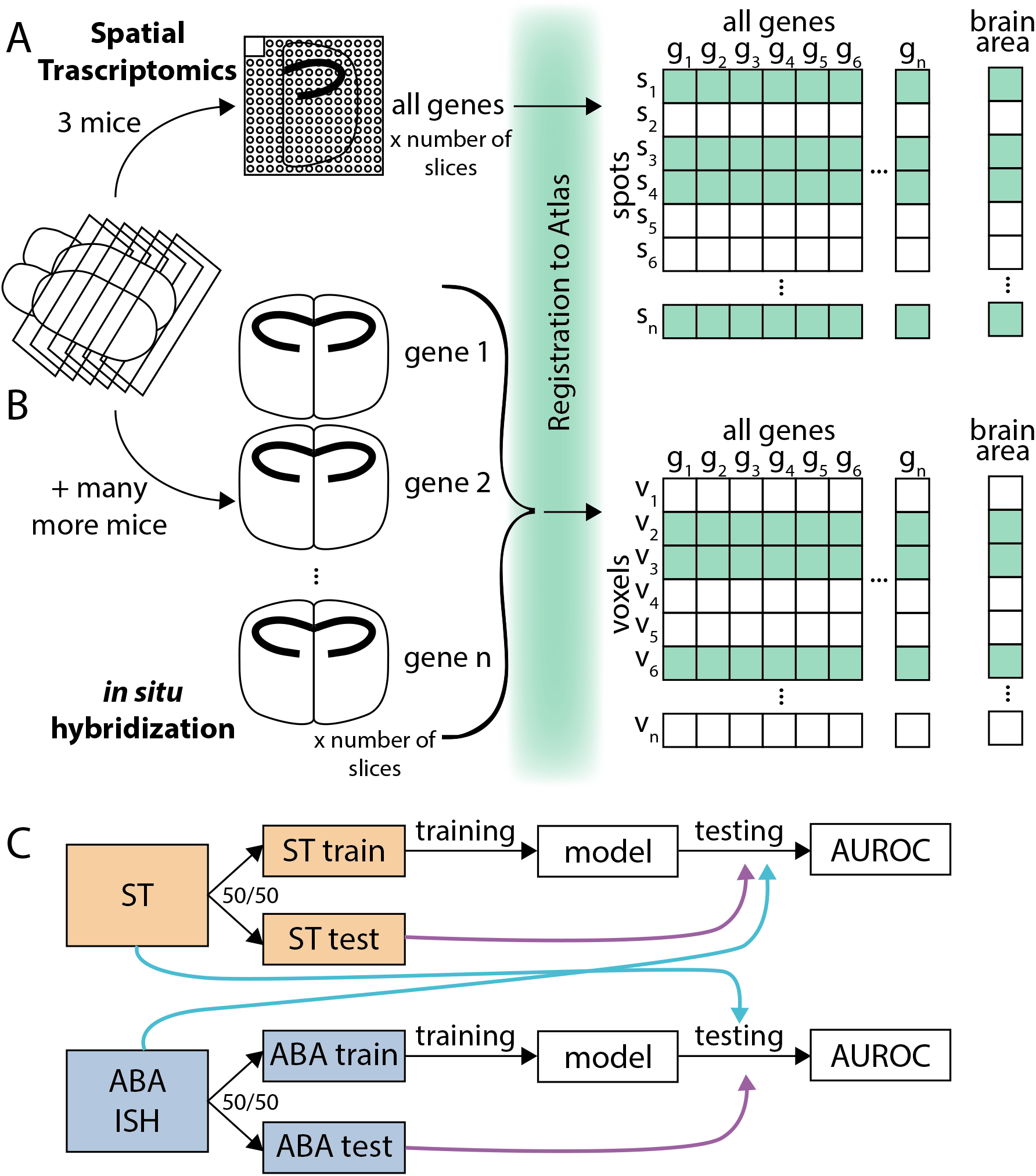
Collection and processing of spatial gene expression datasets. (A) Schematic depicting workflow of collecting whole brain spatial gene expression using Spatial Transcriptomics (ST). Illustration depicts sectioning of mouse brain, tissue from one hemisphere on one Spatial Transcriptomics slide, registration to Allen Reference Atlas, and a layout of the collected data. (B) Schematic depicting workflow of collecting Allen Institute’s whole brain spatial gene expression using *in situ* hybridization (ABA). Illustration depicts similar workflow to (A), but instead of Spatial Transcriptomics capturing all genes in one (three for this dataset) brain, there were many more mice used to collect the whole transcriptome dataset since each brain tissue slice can only be used to probe one gene. (C) Schematic illustrating classification schema. The ST dataset from (A) (orange) and ABA dataset form (B) (blue) were split into 50/50 train/test folds. The training fold was used for model building and the test fold for evaluating the trained model within dataset (purple arrow). Later analysis also applied models trained using the train fold of one dataset to the opposite dataset for testing (light blue arrow).

Using this approach, we show that ARA labels are classifiable using gene expression, but that performance is higher in the ABA than ST. We further demonstrate that models trained in one dataset and tested in the opposite dataset do not reproduce classification performance bidirectionally. We then identify potential biological explanations for the difference in crossdataset performance in classifying brain areas. Finally, we found that although an identifying gene expression profile can always be found for a given brain area, it does not generalize to the opposite dataset. In summary, within each dataset, canonical brain area labels were classifiable and meaningful in gene expression space, but replicability across these two very different assays of gene expression was not robust.

## Results and discussion

### Allen Reference Atlas brain areas are classifiable using gene expression alone

Traditionally, parcellation of the mouse brain has depended on anatomical landmarks and cytoarchitecture, at times, including inter-region connectivity and molecular properties (1,18,19). However, multi-modality agreement of these canonical, atlas brain area subdivisions with gene expression has yet to be assessed at a whole-brain scale. The advent of new technologies for spatial transcriptomics—technologies that preserve the spatial location of RNA transcripts—has enabled the collection of spatially-resolved, whole-transcriptome data in the adult mouse brain to address this question. Specifically, in the present work we ask if canonical brain areas from the Allen Reference Atlas (ARA) (1) are classifiable using two spatial gene expression datasets: the Allen Institute’s own *in situ* hybridization (ABA) data (1,2) and a second dataset collected using Spatial Transcriptomics (ST) (3,17) (Figure 1a, b). Comparing accuracy in classification of ARA brain areas across two technological platforms and datasets allows us to draw conclusions about spatial expression that are more likely to be biological and generalizable than subject to the technical biases of any one dataset.

To determine if we could more generally determine canonical brain areas from spatial gene expression, we first asked if we could do so within each of the two datasets independently. Given the known high correlation structure of gene expression (20), we hypothesized that we could determine the brain area of origin of a gene expression sample using only a subset of the total genes. Fitting these criteria, we chose least absolute shrinkage and selection operator, or LASSO regression (21). LASSO is a regularized linear regression model which minimizes the L1 norm of the coefficients and thus drives most coefficients toward zero. LASSO leaves few genes contributing to the final model and in effect picks “marker genes” of spatial expression in the brain. We use LASSO for two-class classification of all pairwise brain areas in a supervised learning framework with a 50/50 train-test split (Figure 1c) (see methods). The performance of this test set classification is reported using the area under the receiver operating curve (AUROC) (see methods). In this manuscript, we say a brain area pair is classifiable with respect to each other to indicate a high performance in classification with an AUROC greater than 0.5 and generally closer to 1.

After preliminary filtering (see methods), we use this approach in both the ST and ABA to classify all of the smallest brain areas at the leaves of the ARA naming hierarchy against each of the others (461 ST areas; 560 ABA areas), which we will subsequently refer to as leaf brain areas (Figure 1c; see methods). ARA leaf brain areas are classifiable using LASSO (alpha=0.1) from all other leaf brain areas using only gene expression data from (1) the ABA (mean AUROC = 0.996) (Figure 2a, Additional File 1: Supplementary Figure 1a) and from (2) the ST (mean AUROC = 0.883) (Figure 2b, Additional File 1: Supplementary Figure 1b). As expected, performance falls to chance when brain area labels are permuted as a control (ABA mean AUROC = 0.510; ST mean AUROC = 0.501) (Additional File 1: Supplementary Figure 1c-f). Together, these results indicate that in each dataset there is a set of genes whose expression level can be used to identify each of the leaf brain areas from each of the others.

**Figure 2.**
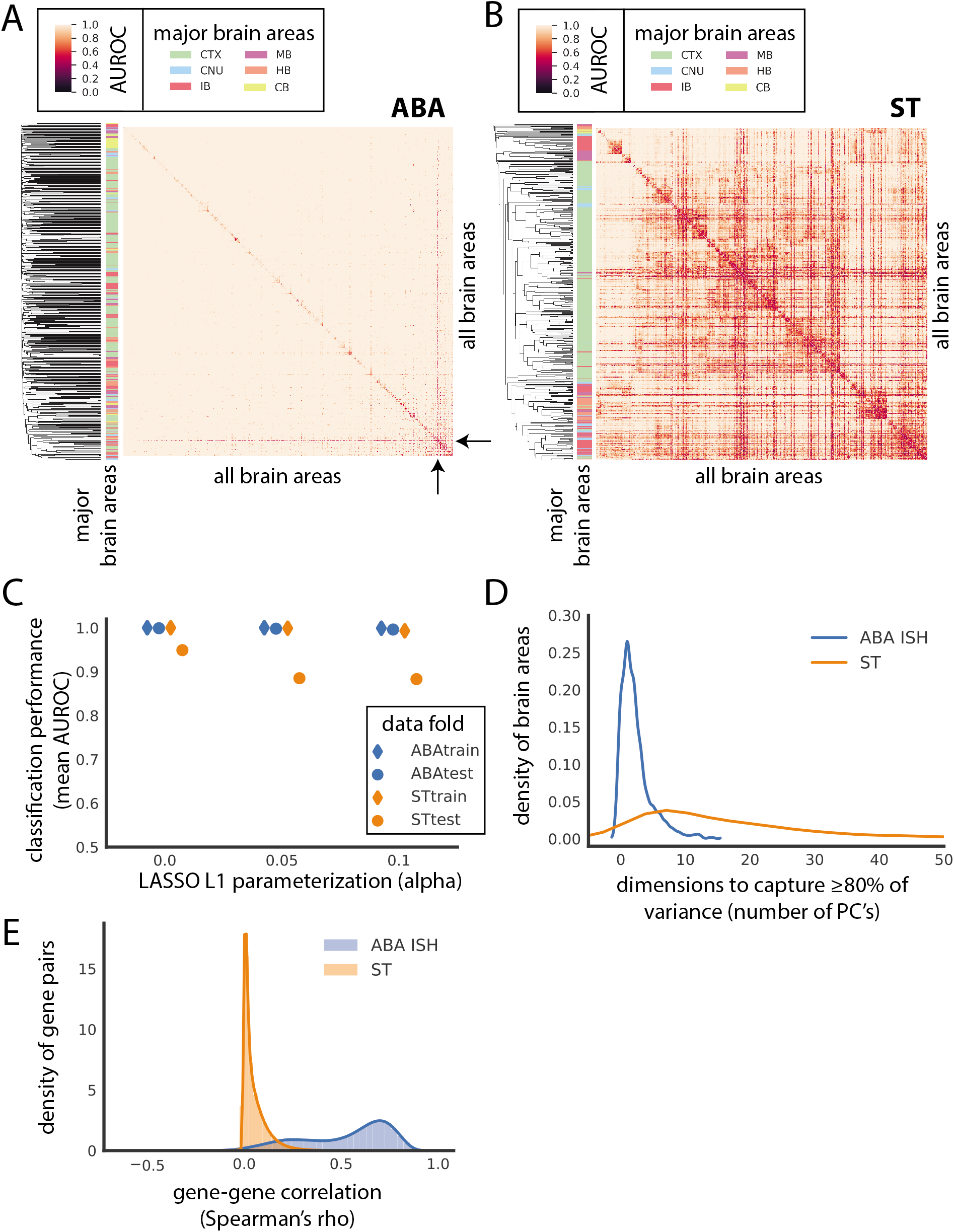
Canonical brain areas are classifiable using gene expression alone in the ABA and ST datasets. Heat map of AUROC for classifying leaf brain areas from all other leaf brain areas in (A) ABA and (B) ST using LASSO (alpha = 0.1). Dendrograms on the far left side represent clustering of leaf brain areas based on the inverse of AUROC; areas with an AUROC near 0.5 get clustered together while areas with an AUROC near 1 are further apart. Color bar on the left represents the major brain structure that the leaf brain area is grouped under. These areas include: cortex (CTX), midbrain (MB), cerebellum (CB), striatum and pallidum (CNU), hindbrain (HB), and thalamus and hypothalamus (IB). (C) Average AUROC (y-axis) of classifying all brain areas from all other brain areas using LASSO across various values of alpha (x-axis): 0, 0.05, and 0.1 for ABA train (blue diamond), ABA test (blue dot), ST train (orange diamond), ST test (orange dot). (D) Number of principal components to capture at least 80% of variance of genes in each of the leaf brain areas after applying PCA to ABA (blue) and ST (orange). ABA brain areas that are larger than ST are randomly down-sampled to have the same number of samples as ST prior to applying PCA. (E) Gene-gene correlations calculated as Spearman’s rho between all pairwise genes across the whole dataset for both the ABA (blue) and ST (orange) independently.

Notably, performance in the ABA is nearly perfect. For the median performing pair of brain areas in ABA (median AUROC = 1), there is a threshold in classification that can be drawn where all instances of one class can be correctly predicted without any false positives (precision = 1). In contrast, in the ST, no such threshold can be found for the median performing (median AUROC = 0.959) brain areas (average precision = 0.846) (see methods). Further, performance in the ABA is consistently higher than the ST across various parameterizations of LASSO (Figure 2c) (see methods). Despite the comparatively lower performance in the ST, clustering brain areas by AUROC shows brain areas belonging to the same major anatomical region grouping together (Figure 2b)(see methods). For example, most brain areas belonging to the cortex group together in the middle of the heat map (green bar on left) with a few interspersed areas. This grouping suggests that patterns of expression track with anatomical labeling. Examining the relative expression of genes that are assayed in both datasets, we see that ranked mean expression is comparable across the two datasets (Spearman’s ρ = 0.599) (Additional File 1: Supplementary Figure 1g) suggesting that the observed difference in performance is not due to lowly expressing genes being highly expressed in the opposite dataset or vice versa.

Observing the nearly perfect performance in the ABA, we next hypothesized that this dataset may be more low-dimensional than suggested by its feature size and may contain many highly correlated features when compared to the ST dataset. Using principal component analysis (PCA), we show that on average 2 PCs are enough to summarize 80% of the variance per brain area in ABA versus 21 PCs in ST (Figure 2d; Additional File 1: Supplementary Figure 1h) (see methods). Further, gene-gene co-expression across the whole dataset is on average higher in the ABA (gene-gene mean Spearman’s rho = 0.525) than the ST (gene-gene mean Spearman’s rho = 0.049) (Figure 2e). The perfect performance, low-dimensionality, and high co-expression all support the idea that while there is meaningful variation in the ABA, it can be captured in one or few dimensions. In summary, canonical ARA brain areas are classifiable from each other using gene expression alone, but performance is likely inflated in the ABA.

An aside of note is that in the ABA the one brain area that is consistently lower performing when classified against most other brain areas is the Caudoputamen (mean AUROC = 0.784) (Figure 2a, black arrows). In the ST, the Caudoputamen is not the lowest performing area, but also has a low mean AUROC (AUROC = 0.619) relative to the other brain areas in ST. In both datasets, the Caudoputamen is the largest leaf brain area composed of the most samples (ABA CP number of voxels = 3012 vs. an average of 85.6 voxels; ST number of spots = 2051 vs. an average of 57 spots). We hypothesized that its relatively larger size could mean that it consists of transcriptomically disparate sub-sections that are not captured with canonical ARA labelling. Falsifying this, we observe that there is no relationship between size and performance in general (Additional File 1: Supplementary Figure 2a,b). Finally, though not an outlier, we do observe that the mean sample correlation for the Caudoputamen in both the ST (mean Pearson’s r = 0.727) and ABA (mean Pearson’s r = 0.665) is slightly lower than the mean in either case (ST mean Pearson’s r = 0.783; ABA mean Pearson’s r = 0.696) (Additional File 1: Supplementary Figure 2c).

### Cross-dataset learning of Allen Reference Atlas brain areas

#### Cross-dataset performance is not bi-directional

Given the low-dimensionality and the near perfect brain area classification performance in the ABA relative to the ST dataset, we hypothesized that the performance of the LASSO models was artificially inflated in the ABA. To explore this hypothesis, we characterized whether LASSO models trained in one dataset would generalize to the opposite dataset (Figure 1c, light blue arrows). For this step we further filtered for (1) 445 leaf brain areas that were represented with a minimum of 5 samples in each dataset and for (2) 14,299 overlapping genes (see methods). LASSO models (alpha = 0.1) trained on ST had a similar within-dataset performance (held out test fold, mean AUROC = 0.884) and cross dataset performance (ABA, mean AUROC = 0.829) (Figure 3a,b), but the reverse is not true. The performance in classifying pairwise leaf brain areas using LASSO models trained in the ABA (held out test fold, mean AUROC = 0.997) falls when testing in the ST (mean AUROC = 0.725) (Figure 3a,c). These results show that the ST dataset is more generalizable to the opposite dataset than the ABA. Additionally, this discrepancy in crossdataset performance suggests that the high performance within the ABA is driven by a property of that dataset not present in the ST.

**Figure 3.**
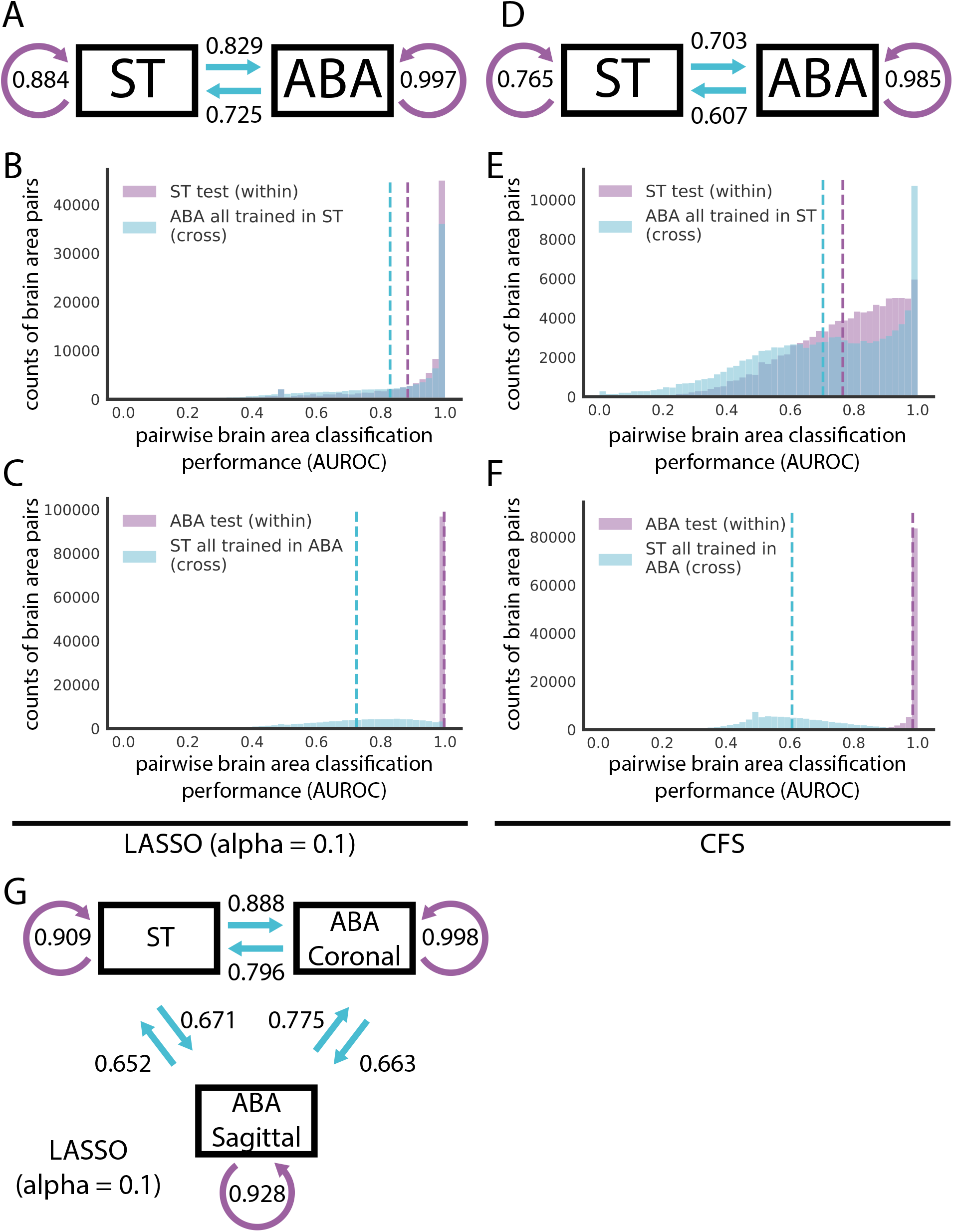
Cross-dataset learning shows that models do not generalize bi-directionally. (A and D) Models trained with overlapping genes and brain areas between ST and ABA datasets are evaluated within dataset on the test fold and across dataset on the entire opposite dataset as illustrated in Figure 1C. Summary diagrams showing mean AUROC for within dataset test set performance (purple arrow) and cross dataset performance with models trained in the opposite dataset (light blue arrow) for (A) LASSO (alpha = 0.1) and (D) CFS. Distributions of AUROCs for within (purple) and cross dataset (light blue) performance for (B) LASSO (alpha = 0.1) trained in ST, (C) LASSO (alpha = 0.1) trained in ABA, (E) CFS trained in ST, and (F) CFS trained in ABA. In all four plots, dashed vertical lines represent the mean of the corresponding colored distribution. (G) Summary diagram showing mean AUROCs using LASSO (alpha = 0.1) for separating out the two planes of slicing in the ABA and treating them alongside the ST dataset as three different datasets for cross-dataset learning. In all three summary diagrams (A, D, G), cross dataset arrows originate from the dataset that the model is trained in and point to the dataset that those models are tested in.

We next ask if the high performance within ABA that is lost when going to ST using models trained in the ABA is specific to the LASSO method or a more general feature of the data. To assess the data more directly, we used to use a second simpler method, correlation-based feature selection (CFS), that eliminates model building and simply picks features that are uncorrelated (22) (see methods). In this way, CFS parallels LASSO which implicitly picks uncorrelated feature sets when minimizing its L1 regularized cost function that penalizes additional features.

Using CFS, we picked 100 randomly seeded feature sets for pairwise comparisons of leaf brain areas (see methods). We then took the single best performing feature set from the train set and evaluated its performance on both the held out test set and cross dataset. We did this in both directions, training on both ST and ABA as with LASSO above. CFS can accurately classify pairwise leaf brain areas in both the ST (test set mean AUROC = 0.765) and in the ABA (test set mean AUROC = 0.985) (Figure 3d-f). As with LASSO, classification in ABA with CFS is on average better performing than in ST. Again following a similar trend as LASSO, the difference in mean cross dataset performance going from ST test set to the ABA (difference in mean AUROC = 0.052; mean ST to ABA cross dataset AUROC = 0.703) is smaller than the reverse (difference in mean AUROC = 0.378; mean ABA to ST AUROC = 0.607) (Figure 3d-f). Altering our analysis approach by averaging the 100 CFS feature sets, we again see a similar pattern in cross dataset performance (ST to ABA difference in mean AUROC = 0.062; ABA to ST difference in mean AUROC = 0.381) (Additional File 1: Supplementary Figure 3a-c). These CFS results indicate that the observed high performance of classification within the ABA and lack of generalization to the ST is not driven by our choice of model. In summary, across both techniques, marker genes can be found to classify pairwise leaf brain areas from each other, but they often do not generalize to the opposite dataset.

#### The sagittal subset of the ABA is the most distinct

With only two datasets it is impossible to distinguish whether the above lack of bidirectionality in cross-dataset learning is driven by (1) the ST being more generalizable or (2) a lack of information in ST that is critical to the high classification performance within ABA. To begin to address this, we took advantage of the separability of the ABA dataset into two distinct datasets: coronal and sagittal. The Allen Institute collected duplicates of many genes; roughly 4,000 genes were collected across both the coronal and sagittal planes of slicing. With these two datasets alongside the ST, we further filtered for 3,737 overlapping genes across the same 445 leaf brain areas (see methods) and computed all pairwise combinations of cross-dataset learning. Notably, using LASSO (alpha = 0.1), training on ST outperforms either plane of ABA in cross dataset predictions: (1) ST to ABA coronal (mean AUROC = 0.888) performs better than ABA sagittal to ABA coronal (mean AUROC = 0.775) and (2) ST to ABA sagittal (mean AUROC = 0.671) performs better than ABA coronal to ABA sagittal (mean AUROC = 0.663)(Figure 3g). Further, the performance of models trained in ABA coronal to ABA sagittal (mean AUROC = 0.663) and ST to ABA sagittal (mean AUROC = 0.671) is lower than that of ABA coronal and ST to each other (ST to ABA coronal mean AUROC = 0.888; ABA coronal to ST mean AUROC = 0.796) (Figure 3g). This shows that the ABA coronal and ST are able to generalize to each other better than to the ABA sagittal. The relative distinctness of the ABA sagittal dataset could be driven by its sparsity-consisting of zeros for more than half of the dataset (53.9%) compared to only 7.5% zeros in the coronal subset. LASSO is able to find a robust set of marker genes within the ABA sagittal that does not reflect the best possible set of genes in the less sparse ABA coronal and ST. Across parametrizations of our model, the sagittal subset of the ABA continues to be the most distinct of the three datasets with the least generalizability (Additional File 1: Supplementary Figure 3d,e).

### Distance in semantic space, but not physical space provides a potential explanation for crossdataset performance

Since the ARA brain areas are organized into a hierarchical tree-like structure based on biology (1), we hypothesized that the semantic distance of any two pairwise brain areas in this tree could provide an explanation for the cross-dataset performance of classifying samples from the same two areas. To investigate this, we used the path length of traversing this tree to get from one brain area to the second area as the measure of distance in the tree (see methods). For the performance of classifying brain areas in both the ST and ABA when trained in the opposite dataset (LASSO, alpha = 0.1), we see an increase in performance (ST to ABA mean AUROC= 0.690 increases to mean AUROC = 0.912; ABA to ST mean AUROC = 0.655 increases to mean AUROC = 0.756) as the semantic distance increases from the minimum value of 2 to the maximum of 15 (Figure 4a,b). As expected, the corresponding increase in performance and semantic distance holds across parameterizations of our linear model (Additional File 1: Supplementary Figure 4a-d). A high AUROC here indicates that the two brain areas are transcriptionally distinct, while an AUROC near 0.5 indicates that they are similar. So, this result implies that distance in semantic space defined by the ARA reflects distance in expression space.

**Figure 4.**
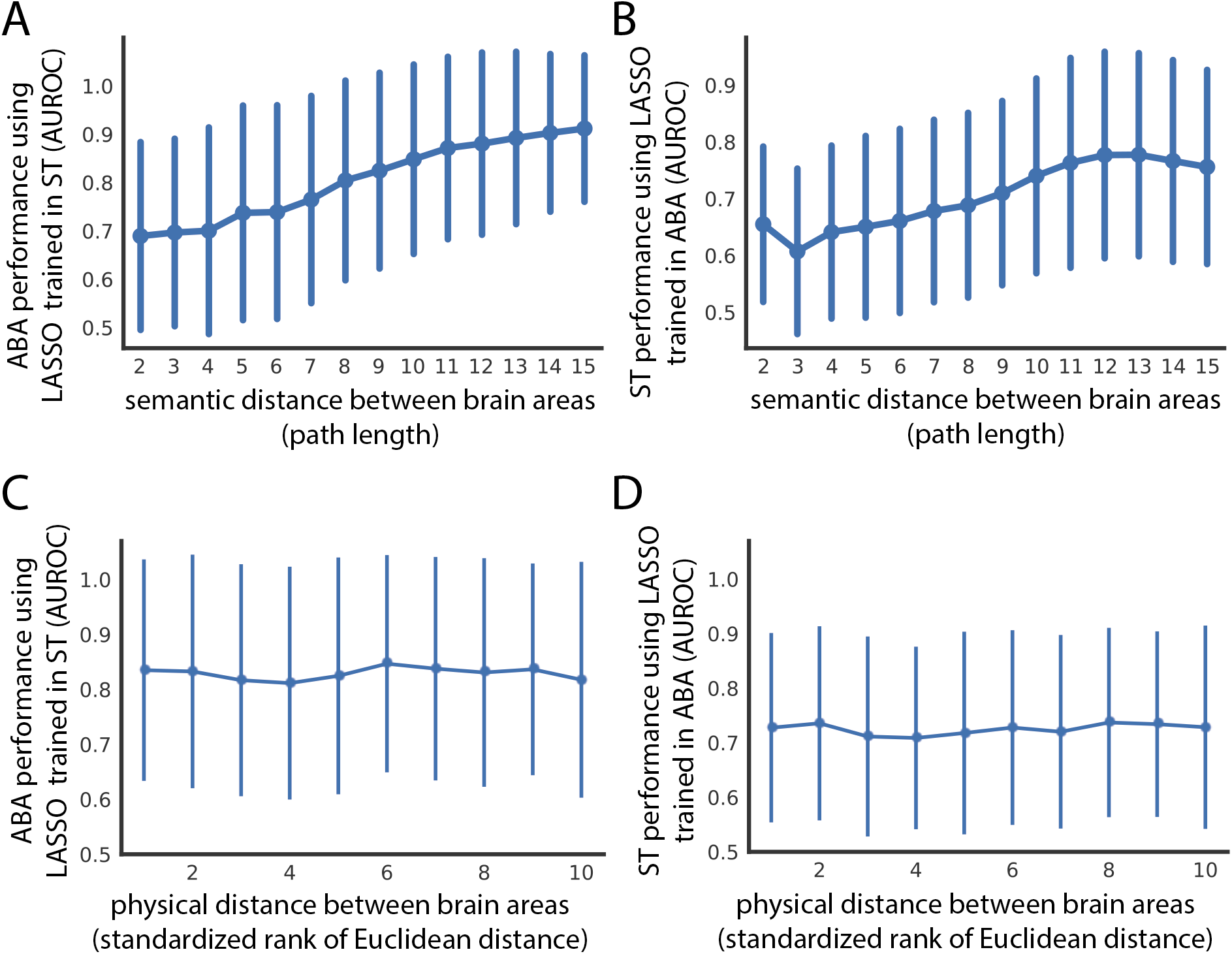
Spatial expression patterns reflect distance in semantic space, but not physical distance in the brain. Cross-dataset AUROCs (x-axis) of classifying all leaf brain areas from all other leaf brain areas for (A) ABA using LASSO (alpha = 0.1) trained in ST and (B) ST using LASSO (alpha = 0.1) trained in ABA as a function of path length (x-axis) in the ARA naming hierarchy between the two brain areas being classified. The same AUROCs (y-axis) from (A) and (B) shown in (C) and (D) respectively as a function of minimum Euclidean distance between the two brain areas in the ARA (x-axis). Euclidean distance on the x-axis is binned into deciles for visualization. All four plots show mean AUROCs (points) with standard deviation (vertical bars).

To further understand the relationship between performance and semantic distance, we next investigated pairs of brain areas with extreme AUROCs at the minimum and maximum semantic distances. We were especially interested in this given the distribution of AUROCs for each distance (Figure 4a, b). Similarly, at the smallest semantic distance of 2, in both ABA and ST trained in the opposite dataset there is a spread in classification performance (Additional File 1: Supplementary Table 1, 2). In both datasets, these brain area pairs involve different cortical layers of the same cortical area. The ARA hierarchy is organized such that within one cortical area, all the layers will have a semantic distance of 2 between each other. So, a pair of brain areas with a high AUROC and semantic distance of 2 often involves two non-neighboring layers of a cortical area (i.e. primary auditory cortex layer 6b and layer 4 in ST trained in ABA) (Additional File 1: Supplementary Table 2). This trend is in line with our expectation as cortical layers are known to have distinct expression profiles driven in part by distinct cell types (23–26). Alternatively, a pair of brain areas with an AUROC near 0.5 and a semantic distance of 2 can involve two neighboring layers of a cortical area (i.e. primary visual area layer 6a and layer 6b in ST trained in ABA) (Additional File 1: Supplementary Table 2). This too is not surprising because, despite distinctness in cortical layer expression, we expect some overlap between physically neighboring areas in terms of expression profiles due to errors introduced in sampling and in registration to the reference atlas. This example illustrates one way in which semantic distance is not synonymous to physical distance.

Since semantic distance does not perfectly capture the actual distance between brain areas, we next looked at classification performance as a function of physical distance directly. Specifically, we asked: Is performance in classifying pairwise leaf brain areas cross-dataset being driven by physical proximity/distance alone? This allows us to determine if differential gene expression patterns between brain areas are simply due to spatial differences in gene expression. Cross-dataset performance was examined with respect to the minimum Euclidean distance between the two brain areas in the ARA (see methods). There is no trend between physical distance and AUROCs from either cross-dataset assessment using LASSO models trained in the opposite dataset (ABA to ST Pearson’s r = −0.026; ST to ABA Pearson’s r = 0.056) with the mean performance remaining similar at the minimum (ABA to ST mean AUROC = 0.651; ST to ABA mean AUROC = 0.702) and maximum distance (ABA to ST mean AUROC = 0.697, change in AUROC = +0.046; ST to ABA mean AUROC = 0.690, change in AUROC = −0.012) (Figure 4c, d) (see methods). Across model parameterizations, there is similarly no relationship between distance and performance (Additional File 1: Supplementary Figure 4e-h). This result alongside the positive relationship seen between performance and semantic distance, implies that canonical brain area labels are meaningful and provide more information than physical distance alone in the space of gene expression.

### Finding a uniquely identifying gene expression profile for individual brain areas

#### Within one dataset a gene expression profile can uniquely identify one brain area, but it does not generalize to the opposite dataset

Thus far, we have focused on the classification of leaf brain areas from other leaf brain areas. However, this does not determine if we can uniquely identify a given brain area from the whole brain using gene expression. To tackle this, we trained linear models for one leaf brain area against the rest of the brain (one versus all) and tested that same model’s performance in classifying the same leaf brain area against all others (one versus one across all leaf brain areas) (Figure 5a). Unfortunately, for most leaf brain areas, LASSO fails to fit a model with very light regularization (alpha = 0.01) to classify it against the rest of the brain in both the ST (mean train AUROC = 0.554) and the ABA (mean train AUROC = 0.593) (Additional File 1: Supplementary Figure 5a-d). The few leaf brain areas that are able to be classified from the rest of the brain using LASSO have a nearly identical performance in the one versus all case as in testing against all other leaf brain areas (Figure 5b; Additional File 1: Supplementary Figure 5a, b). At a higher regularization weight (alpha = 0.05), most one versus all models fail to be trained (ST mean train AUROC = 0.501; ABA mean train AUROC = 0.502) (Figure 5b; Additional File 1: Supplementary Figure 5e-h). Due to this inability to train a model in the one versus all case for most leaf brain areas, we turned to no regularization of LASSO (alpha = 0), which is equivalent to linear regression. Using linear regression, performance of models fit in the one versus all case correlates nearly perfectly with the average performance of the same model in one versus one. This nearly identical performance is true in both the ST (mean distance from identity line = 0.005) and ABA datasets (mean distance from identity line = 0.001) (Figure 5b-d) (see methods). This result demonstrates that within a dataset, we can find an identifying gene expression profile of a brain area that uniquely identifies it.

**Figure 5.**
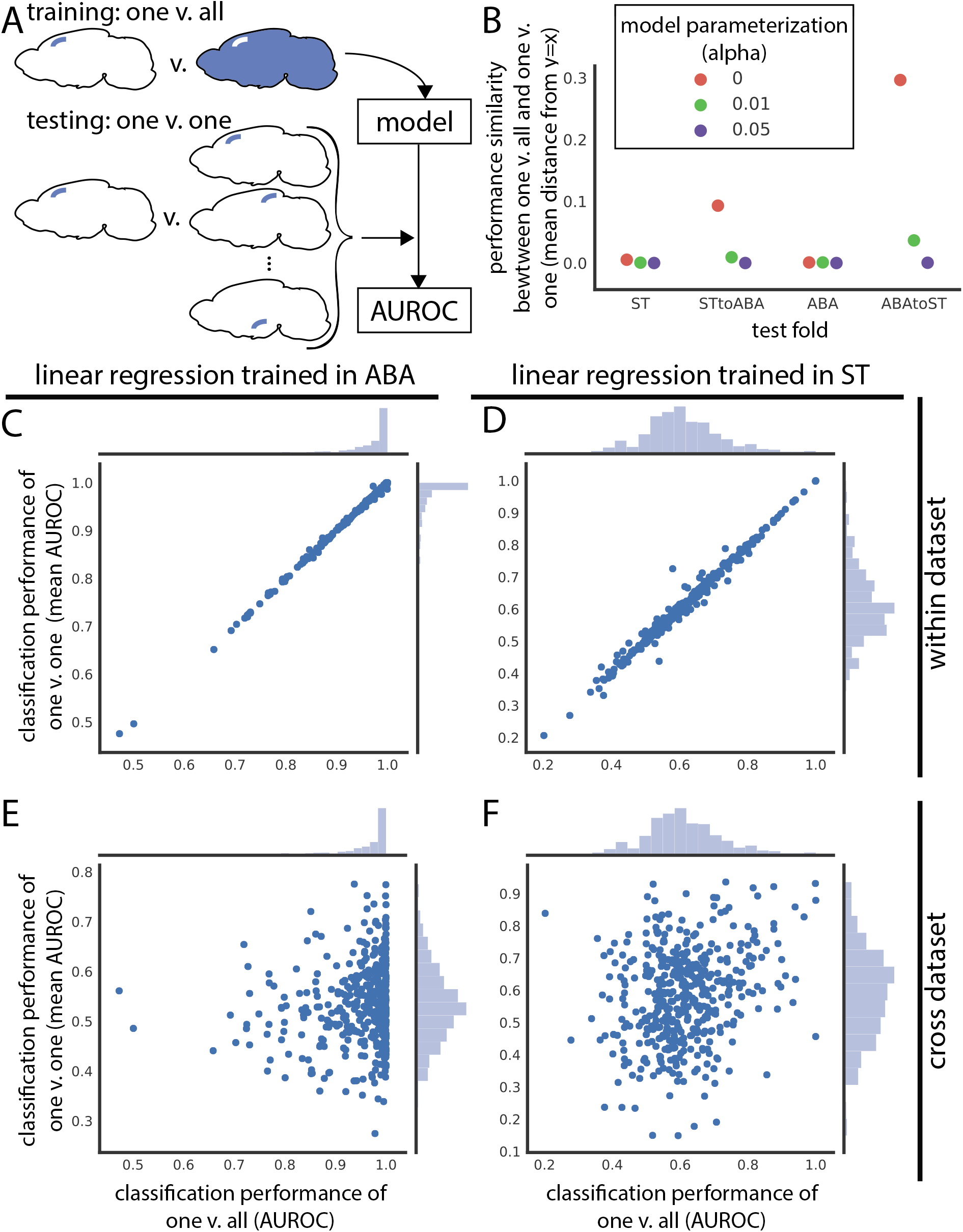
Leaf brain area expression profiles are identifiable within dataset, but do not generalize cross dataset. (A) Schematic depicting training and testing schema for panels in this figure. Models are trained to classify one leaf brain area against the rest of the brain (one v. all) and then used to test classification of that brain area against all other leaf brain areas (one v. one) within and across dataset. (B) Performance similarity between one v. all and one v. one reported as the mean absolute value of distance from identity line for scatter plots of testing in one v. all against one v. one. Performance similarity shown for within ST, ST to ABA, within ABA, and ABA to ST across linear regression (red), LASSO (alpha = 0.01) (green), and LASSO (alpha = 0.05) (violet). Linear regression one v. all test set performance (x-axis) versus average one v. one performance of the same model (y-axis) in (C) ABA and (D) ST. (E) Assessment of the same ABA one v. all linear regression model (x-axis) in one v. one classification in the ST dataset (y-axis). (F) Same as (E), but one v. all linear regression trained in ST (x-axis) and one v. one classification of these models in ABA (y-axis).

Since we could robustly identify a gene expression profile to identify a brain area within one dataset, we next asked if these profiles can generalize to the opposite dataset. Using the same models trained in one versus all in either the ST or ABA, we classified the same brain area against all other brain areas (one versus one) in the second dataset. The one versus all trained linear models (alpha = 0) do not generalize cross-dataset for either ABA to ST (mean distance from identity line = 0.296) or the reverse (ST to ABA mean distance from identity line = 0.093) (Figure 5b, e-f). This lack of cross dataset performance similarly holds for other parameterizations (Figure 5b; Additional File 1: Supplementary Figure 5c, d, g, h). The identifying gene expression profile of a leaf brain area is not generalizable to a new dataset that is not used in defining that profile. So, while we can uniquely identify a brain area using gene expression within one dataset, that identification profile does not extend to the second dataset.

### Conclusions

Across disciplines, benchmarking studies have helped to advance their respective fields and set standards for future research (16,27–29). Neuroscience is no exception. Given the complexity of the brain, however, the addition of multimodal information is especially coveted. Studying the brain from a variety of perspectives, across data modalities, technology platforms, and experiments, can give a complete, composite understanding of its biology (30). With this in mind, we linked two modalities to ask: can we capture canonical, anatomically-defined brain areas from the Allen Reference Atlas using spatial gene expression alone? And, how well does this replicate across two transcriptomic datasets collected using different platforms?

Principally, we showed that ARA brain labels are classifiable using only gene expression, but highlighted a lack of generalizability across spatial transcriptomic datasets. Within datasets, we are able to distinguish brain areas from each other with high performance. We were further able to uniquely identify a brain area within dataset; training on one brain area against the entire rest of the brain generalizes to testing on that same brain area against all other leaf brain areas. Notably, within dataset performance was on average higher in the ABA than the ST, which led to a lack of cross dataset generalizability when training in the ABA and testing in ST; this phenomenon was not present in reverse. However, in both cases, there is an observed trend linking an increase in mean cross dataset performance with increased semantic distance in the ARA brain area label organization. There was no link in performance when compared to physical distance in the brain suggesting ARA labels are meaningful in expression space and we are not simply detecting spatial differences in gene expression.

It is important to point out that our benchmarking study at its core only involves two independent datasets, although both are extraordinary in scope. Limited to two datasets, it is impossible to tell whether one dataset or the other is closer to representing the ground truth. Further, the ABA and ST datasets use two fundamentally different techniques: the ABA data reports average pixel intensity from *in situ* hybridization and the ST approach is an RNA capture technique followed by sequencing that reports read counts. One likely source of discrepancy between the ST and ABA datasets is downstream processing. After the collection of the raw expression data, each of the two datasets also undergoes a unique registration step to the ARA. The ABA uses an iterative approach that involves registration of the 3D brain volume with interspersed smoothing steps (2), while the ST dataset is registered on a slice by slice basis to the nearest representative 2-D ARA slice using anatomical landmarks (31). Beyond registration, there are additional concerns about the stability of the ARA brain area labels since there are inconsistencies with other brain atlases and even across versions of the ARA (32–34). Brain atlases are an imperfect formalism of brain substructure, but are the best systematic representation to test spatial gene expression by biological areas.

As types and prevalence of spatial gene expression approaches continue to increase (6), whole-brain spatial gene expression datasets will follow. By continuing to integrate these emerging datasets, we will be able to perform more robust meta-analysis, giving us a deeper understanding of spatial gene expression with respect to ARA labels. An added benefit of the continued incorporation of additional datasets, is that at some point, differences in experimental platforms and registration approaches will only contribute to the robustness of any generalizability claims. With the interpretable cross-dataset analysis framework established here, we have laid a foundation for the quantification of generalizability of future whole-brain spatial datasets. We hope this work will serve to usher forward the next wave of meta-analysis of spatial gene expression in the adult mouse brain as a route forward toward integration of distinct data types – location and expression – to form the beginnings of a robust, multi-modality understanding of the mammalian brain.

## Methods

### Spatial Transcriptomics Data (ST)

Spatial Transcriptomics (ST) is an array-based approach where a tissue section is placed on a chip containing poly-T RNA probes that the mRNA transcripts present in the tissue can hybridize to (3). These probes tile the chip in 100 micron diameter spots and contain barcodes specific to that spot so that RNA sequencing reads can be mapped back to their original grid location. Note that the probe spots are not perfectly adjacent to each other, but have a center to center distance of 200μm (3).

Here, we used a previously published spatial gene expression dataset containing 75 coronal slices from one hemisphere of the adult mouse brain across three animals (17). The coronal slices were mapped to the Allen Mouse Brain Reference Atlas using a non-rigid transformation approach (31). In total, this dataset contains 34,103 ST spots across 23,371 genes (17).

### Allen Brain Atlas in situ hybridization Data (ABA)

The Allen Brain Atlas (ABA) adult mouse in situ hybridization (ISH) dataset consists of a transcriptome-wide assay of expression in inbred WT mice using single molecule ISH (1). To assay the whole transcriptome, many WT mouse brains were sliced into 25μm thick slices containing 8 interlayered sets for subsequent single molecule hybridization or for staining to create the reference atlas. This results in a z resolution of 200μm for each gene. These independent image series are subsequently reconstructed to three dimensions and registered to the reference brain atlas in interlayered steps (2). There are 26,078 series, or experiments, across both coronal and sagittal planes with 19,942 unique genes represented. These 3D registered reconstructions are then segmented to 200μm^3^ voxels with an associated brain area label. There are 159,326 voxels, with 62,529 mapping to the brain. Gene expression for each of the assayed genes was quantified in these voxels from the imaged data as energy values which is defined as the sum of expression pixel intensity divided by the sum of all pixels.

The quantified ISH energy values dataset was downloaded from the Allen Brain Atlas website (http://help.brain-map.org/display/mousebrain/API) through their API on March 12, 2019.

### Allen Institute reference brain ontology, leaf brain areas, and path length

The Allen Institute reference brain atlas has organized brain areas into a hierarchy described by a tree data structure. Leaf brain areas are defined here as brain areas that constitute leaves on the ontology tree, i.e. they have no children. Leaf brain areas represent the most fine-scale parcellation of the brain. Using leaf brain areas circumvents the fact that the depth of the tree representing the hierarchical naming structure of brain areas in the ARA is not uniform. Path length refers to the number of steps required to go from one brain area to another in this tree.

### Data filtering and train/test split

The ST data was pre-processed to remove ST spots mapping to ambiguous regions, fiber tracts, or ventricular systems and to remove genes that were expressed in less than 0.1% of samples. This left 30,780 ST spots, or samples, with 16,557 genes. For within ST analyses, this dataset was further filtered to 461 leaf brain areas that each had a minimum of 5 spots. In all analyses these spots are subsequently randomly split into train and test sets with a 50/50 split. The train/test split is random, but stratified for brain areas so that each fold has roughly 50% of the samples belonging to each brain area. For within ST training and testing, n-fold (here, 2) cross validation was used and results are reported as a mean across folds.

Similarly, the ABA data was filtered for only voxels mapping to the reference brain and genes with expression in at least 0.1% of samples. This gives 26,008 series across 62,527 voxels also split as described for ST into 50/50 train and test folds. There are 4,972 genes that are assayed more than once across independent experimental series. Except for the analyses separating out the two planes of the ABA data (detailed below), genes duplicated across series were averaged for each voxel. For within ABA analyses, this dataset was further filtered to 560 leaf brain areas that each had a minimum of 5 voxels prior to the train/test split. As with ST, within ABA training and testing, n-fold (here, 2) cross validation was used and results are reported as a mean across folds.

For cross-dataset learning, both datasets were further filtered for 445 leaf brain areas that were represented with a minimum of 5 samples in each dataset. Genes were also filtered for those present in both datasets resulting in 14,299 overlapping genes between the two. For analyses separating out the two ABA planes, a similar mapping process was used to determine overlaps between each of the planes and the ST data. This resulted in 3737 overlapping genes across the same 445 leaf brain areas. Genes that were duplicated in the ABA dataset with independent imaging series within a plane were averaged.

### Area under the receiver operating characteristic (AUROC), clustering using AUROC, and precision

The AUROC is typically thought of as calculating the area under the curve of true positive rate as a function of false positive rate. Many uses of AUROC report the estimated area under this curve. Here, the analytical AUROC is calculated since it is both computationally tractable and accurate. It is given by

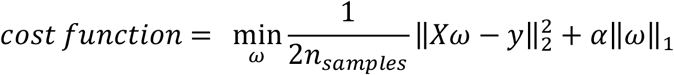

where ranks are the ranks of each positive label sorted by feature and N_Pos_ and N_Neg_ are the number of positive and negative labels respectively. An AUROC of 0.5 indicates that the task being evaluated is performing at chance, while an AUROC of 1 indicates perfect performance. Any AUROCs of 0 were removed from downstream reporting of distributions and mean AUROCs. For within dataset analyses, AUROC’s are reported as the mean across 2-fold cross validation.

Clustering by AUROC is done by converting AUROC to a similarity metric by subtracting 0.5 to center the AUROC values at 0.5 and taking the absolute value. The rationale is that if a classification task performs with an AUROC of 0.5, the two classes are so similar that they are not distinguishable so they should be grouped closely.

Here, we calculate precision for the median performing brain area pair given by AUROC for within dataset analysis. We use a threshold that includes all instances of the positive label, here, all instances of one brain area. Precision is calculated as:

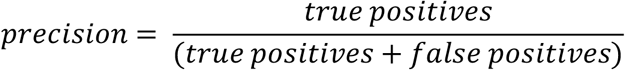

Note, that for while we report the median performing brain area pairs as the mean AUROCS across 2 folds, we investigate the precision using the median of each fold independently and then average the precision of all median brain area pairs.

### LASSO

Least absolute shrinkage and selection operator, or LASSO regression uses a L1 penalty for fitting the linear regression model (21). The cost function to minimize is given by:

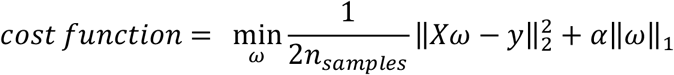

where X represents the matrix of feature values, y the target values, ω the coefficients, and α the constant value with which to weight the regularization. A small α gives little regularization (α = 0 is equivalent to regular linear regression). An L1 penalty minimizes the absolute value of coefficients, which has an effect of pushing many coefficients toward zero. This is beneficial for highly correlated data to find an optimal set of features among correlated genes, or features, to use for prediction.

In this manuscript, LASSO models are fit using coordinate descent according to the scikit-learn library (35). The hyperparameter α was set as noted in the main text. For additional details on parameterization, see code scripts (repository availability below).

### Assessing dimensionality of data using Principal Component Analysis (PCA)

Principal Component Analysis (PCA) as implemented in scikit-learn (35), was used to determine the dimensionality of both datasets. PCA was applied to the genes, or features, for each leaf brain area in the ABA and ST datasets. The total number of components to use for dimensionality reduction was set to be equal to the number of samples in each area. ABA areas were down-sampled to have the same number of samples as the corresponding brain area in ST. There are 77 brain areas that exceptionally have fewer samples in ABA than ST, so when downsampling ABA for these 77 areas, the original sample size was used. Dimensionality of the brain areas were than accessed as the number of PCs needed to explain at least 80% of the variance.

### Differential expression and correlation-based feature selection (CFS)

Differential expression (DE) in genes is assayed using Mann-Whitney U (MWU). Resulting p-values are not corrected for multiple hypothesis testing since p-values are only used to threshold for very extreme DE genes across brain area comparisons. The un-corrected p-values themselves are not reported as a measure for significant DE.

Correlation-based feature selection (CFS) is a feature selection technique that explicitly picks uncorrelated features (22). Here, a greedy approach to CFS was implemented. The algorithm first choses a random seed or gene within the top 500 differentially expressed genes. The next gene is then chosen as the lowest correlated gene to the first one and kept if the set AUROC improves. Subsequent genes are chosen as the least correlated on average to the genes already in the feature set. The algorithm stops once the AUROC is no longer improving. The final set of genes chosen using CFS are then aggregated by equally by averaging the values of all chosen genes for each sample. For more details on exact implementation see code scripts (repository available below).

For the cross-dataset analysis, when un-specified, 100 feature sets were chosen using this approach and the single best performing feature set was then evaluated in both the within dataset test set and the cross dataset test set. When indicated accordingly, the 100 CFS feature sets were averaged instead of reporting the performance of the best set alone.

### Euclidean distance between two brain areas

In addition to brain area labels, the ABA dataset contains x, y, z coordinates for each voxel in the ARA space. So, physical distance between two brain areas is calculated as the Euclidean distance between the two closest voxels where each voxel belongs to one or the other brain area. Due to the symmetry of brain hemispheres, distance was only calculated in one hemisphere by filtering for voxels with a z-coordinate less than 30. This z-coordinate was visually determined to be the midline of the brain based on 3-D visualization of the voxel coordinates. Euclidean distances between brain areas calculated in this manner were used for both the ST and ABA datasets since both are registered to the ARA.

### Mean distance from identity line

To assess the replicability of models trained in one brain area versus the rest of the brain (one versus all) in classifying that same brain area against all the others (one versus one), the mean absolute Euclidean distance of a scatter plot of those two values from the identity line was calculated. This was done to assess how similar the values are in the one versus all case are to the one versus one case for each pair of brain areas. Correlation was found to be lacking because it could yield high correlations when the one versus all and one versus one values were quite different for a given point.

## Availability of data and materials

All code used for the analyses described in this manuscript was written in Python 3.7 with supporting packages: jupyterlab 1.0.9, h5py 2.9.0, numpy 1.16.4, scipy 1.3.1, pandas 0.25.0, scikit-learn 0.21.2, matplotlib 3.1.0, and seaborn 0.9.0. All Jupyter notebooks and scripts are available on GitHub at www.github.com/shainalu/spatial_rep.

The ABA data is available publicly for download directly from the Allen Brain Atlas website (http://help.brain-map.org/display/api/Allen%2BBrain%2BAtlas%2BAPI). ST data (from Ortiz *et al*., 2020) is available at molecularatlas.org.

## Author contributions

AZ and JG conceived the study. SL, AZ, and JG designed the analyses and computational experiments with input from SF, DF, and KM. SL performed the analyses. CO pre-processed and provided the ST data under guidance from KM. SL wrote the manuscript with feedback from SF, JG, and AZ. All authors read and approved the final manuscript.

## Acknowledgements

The authors would like to thank Leon French for providing insight on systematically accessing the Allen Institute data, Manthan Shah for executing this, and Nathan Fox for streamlining datasets used. The authors would also like to thank Sara Ballouz, Xiaoyin Chen, Megan Crow, Aki Funamizu, Benjamin Harris, Longwen Huang, Risa Kawaguchi, Elyse Schetty, Colin Stoneking, and Alex Vaughan for useful input and discussion.

## References

1. Lein ES, Hawrylycz MJ, Ao N, Ayres M, Bensinger A, Bernard A, et al. Genome-wide atlas of gene expression in the adult mouse brain. Nature. 2007 Jan 6;445(7124):168–76.

2. Ng L, Pathak SD, Kuan C, Lau C, Dong H, Sodt A, et al. Neuroinformatics for genome-wide 3D gene expression mapping in the mouse brain. IEEE/ACM Trans Comput Biol Bioinforma. 2007;4(3):382–92.

3. Ståhl PL, Salmén F, Vickovic S, Lundmark A, Navarro JF, Magnusson J, et al. Visualization and analysis of gene expression in tissue sections by spatial transcriptomics. Science. 2016 Jul 1;353(6294):78–82.

4. Rodriques SG, Stickels RR, Goeva A, Martin CA, Murray E, Vanderburg CR, et al. Slide-seq: A scalable technology for measuring genome-wide expression at high spatial resolution. Science. 2019 Mar 29;363(6434):1463–7.

5. Vickovic S, Eraslan G, Salmén F, Klughammer J, Stenbeck L, Schapiro D, et al. High-definition spatial transcriptomics for in situ tissue profiling. Nat Methods. 2019 Oct 1;16(10):987–90.

6. Asp M, Bergenstråhle J, Lundeberg J. Spatially Resolved Transcriptomes—Next Generation Tools for Tissue Exploration. BioEssays. 2020 May 4;1900221.

7. Economo MN, Viswanathan S, Tasic B, Bas E, Winnubst J, Menon V, et al. Distinct descending motor cortex pathways and their roles in movement. Nature. 2018;563(7729):79–84.

8. Bendesky A, Kwon YM, Lassance JM, Lewarch CL, Yao S, Peterson BK, et al. The genetic basis of parental care evolution in monogamous mice. Nature. 2017 Apr 27;544(7651):434–9.

9. Moffitt JR, Bambah-Mukku D, Eichhorn SW, Vaughn E, Shekhar K, Perez JD, et al. Molecular, spatial, and functional single-cell profiling of the hypothalamic preoptic region. Science. 2018;362(6416).

10. Cadwell CR, Palasantza A, Jiang X, Berens P, Deng Q, Yilmaz M, et al. Electrophysiological, transcriptomic and morphologic profiling of single neurons using Patch-seq. Nat Biotechnol. 2016;34(2):199–203.

11. Hanchate NK, Lee EJ, Ellis A, Kondoh K, Kuang D, Basom R, et al. Connect-seq to superimpose molecular on anatomical neural circuit maps. Proc Natl Acad Sci U S A. 2020;117(8):4375–84.

12. Chen X, Sun YC, Zhan H, Kebschull JM, Fischer S, Matho K, et al. High-Throughput Mapping of Long-Range Neuronal Projection Using In Situ Sequencing. Cell. 2019;179(3):772–786.e19.

13. Huang L, Kebschull JM, Fürth D, Musall S, Kaufman MT, Churchland AK, et al. BRICseq Bridges Brain-wide Interregional Connectivity to Neural Activity and Gene Expression in Single Animals. Cell. 2020;182(1):177–188.e27.

14. Sun Y-C, Chen X, Fischer S, Lu S, Gillis J, Zador AM. Integrating barcoded neuroanatomy with spatial transcriptional profiling reveals cadherin correlates of projections shared across the cortex. bioRxiv. 2020;2020.08.25.266460.

15. Canales RD, Luo Y, Willey JC, Austermiller B, Barbacioru CC, Boysen C, et al. Evaluation of DNA microarray results with quantitative gene expression platforms. Vol. 24, Nature Biotechnology. 2006;p. 1115–22.

16. Shi L, Reid LH, Jones WD, Shippy R, Warrington JA, Baker SC, et al. The MicroArray Quality Control (MAQC) project shows inter- and intraplatform reproducibility of gene expression measurements. Nat Biotechnol. 2006;24(9):1151–61.

17. Ortiz C, Navarro JF, Jurek A, Märtin A, Lundeberg J, Meletis K. Molecular atlas of the adult mouse brain. Sci Adv. 2020;6(26):eabb3446.

18. Crick F, Jones E. Backwardness of human neuroanatomy. Nature. 1993;361(6408):109–10.

19. MacKenzie-Graham A, Lee EF, Dinov ID, Bota M, Shattuck DW, Ruffins S, et al. A multimodal, multidimensional atlas of the C57BL/6J mouse brain. J Anat. 2004;204(2):93–102.

20. Eisen MB, Spellman PT, Brown PO, Botstein D. Cluster analysis and display of genomewide expression patterns. Proc Natl Acad Sci. 1998;95(25):14863–8.

21. Tibshirani R. Regression shrinkage and selection via the LASSO. J R Stat Soc B. 1996;58(1):267–88.

22. Hall MA. Correlation-based Feature Selection for Machine Learning. The University of Waikato; 1999.

23. Yao Z, Liu H, Xie F, Fischer S, Adkins R, Aldrige A, et al. An integrated transcriptomic and epigenomic atlas of mouse primary motor cortex cell types. bioRxiv. 2020;2020.02.29.970558.

24. Yao Z, Nguyen TN, van Velthoven C, Goldy J, Sedeno-Cortes A, Baftizadeh F, et al. A taxonomy of transcriptomic cell types across the isocortex and hippocampal formation. bioRxiv. 2020;2020.03.30.015214.

25. Codeluppi S, Borm LE, Zeisel A, La Manno G, van Lunteren JA, Svensson CI, et al. Spatial organization of the somatosensory cortex revealed by osmFISH. Nat Methods. 2018;15(11):932–5.

26. Poulin JF, Tasic B, Hjerling-Leffler J, Trimarchi JM, Awatramani R. Disentangling neural cell diversity using single-cell transcriptomics. Vol. 19, Nature Neuroscience. Nature Publishing Group; 2016. p. 1131–41.

27. Su Z, Łabaj PP, Li S, Thierry-Mieg J, Thierry-Mieg D, Shi W, et al. A comprehensive assessment of RNA-seq accuracy, reproducibility and information content by the Sequencing Quality Control Consortium. Nat Biotechnol. 2014 Sep 1;32(9):903–14.

28. Meta-analysis in basic biology. Vol. 13, Nature Methods. Nature Publishing Group; 2016. p. 959.

29. Robinson MD, Vitek O. Benchmarking comes of age. Vol. 20, Genome Biology. BioMed Central Ltd.; 2019. p. 205.

30. Lein E, Borm LE, Linnarsson S. The promise of spatial transcriptomics for neuroscience in the era of molecular cell typing. Science. 2017;358(6359):64–9.

31. Fürth D, Vaissière T, Tzortzi O, Xuan Y, Märtin A, Lazaridis I, et al. An interactive framework for whole-brain maps at cellular resolution. Nat Neurosci. 2018;21(1):139–49.

32. Azimi N, Yadollahikhales G, Argenti JP, Cunningham MG. Discrepancies in stereotaxic coordinate publications and improving precision using an animal-specific atlas. Vol. 284, Journal of Neuroscience Methods. Elsevier B.V.; 2017. p. 15–20.

33. Chon U, Vanselow DJ, Cheng KC, Kim Y. Enhanced and unified anatomical labeling for a common mouse brain atlas. Nat Commun. 2019;10(1):1–12.

34. Wang Q, Ding SL, Li Y, Royall J, Feng D, Lesnar P, et al. The Allen Mouse Brain Common Coordinate Framework: A 3D Reference Atlas. Cell. 2020;181(4):936–953.e20.

35. Pedregosa F, Michel V, Grisel O, Blondel M, Prettenhofer P, Weiss R, et al. Scikit-learn: Machine Learning in Python. Vol. 12, Journal of Machine Learning Research. 2011.

