## Additional File 1: Supplementary Figures 1-5, Supplementary Tables 1-2 for "Replicability of spatial gene expression atlas data from the adult mouse brain"

**Supplementary Information for "Replicability of spatial gene expression atlas data from the adult mouse brain"**

**Table of Contents**


| | |
|---|---|
| Supplementary Figure and Table Legends | 1-4 |
| Supplementary Figures | 5-9 |
| Supplementary Tables | 10-14 |


**Supplementary Figure Legends**

**Supplementary Figure 1.**

Histogram of classification performance of LASSO (alpha = 0.1) in (A) ABA test fold and (B) ST. (A) and (B) represent the upper triangular of (Figure 2A) and (Figure 2B) respectively. Black dashed vertical line represents the mean.

Upper triangular (C) histogram and (E) heatmap map of AUROC for classifying leaf brain areas from all other leaf brain areas in ABA using LASSO (alpha = 0.1) when brain area labels are randomly permuted. (D) and (F) same as (C) and (E) respectively, but for ST with labels permuted. For heatmaps (E-F), dendrograms on the far left side represent clustering of leaf brain areas based on the inverse of AUROC; areas with an AUROC near 0.5 get clustered together while areas with an AUROC near 1 are further apart. Color bar on the left represents the major brain structure that the leaf brain area is grouped under. These areas include: cortex (CTX), midbrain (MB), cerebellum (CB), striatum and pallidum (CNU), hindbrain (HB), and thalamus and hypothalamus (IB).

(G) Relative expression between the ST and ABA datasets as a density plot. Expression is plotted as the ranked mean for each gene across all samples.

(H) Cumulative explained variance curves for PCA in ST (orange) and ABA (blue). Each curve represents one leaf brain area. The total number of principal components per brain area is equal to the number of samples in that area. ABA areas that had more samples than ST are randomly down-sampled accordingly. For visualization, the Caudopoutamen is not shown for either dataset.

**Supplementary Figure 2.**

Test set AUROC in (A) ST and (B) ABA as a function of the number of samples per brain area. The minimum of the two brain areas involved in classification is shown.

(C) Distribution of sample correlation within each of the leaf brain areas for ABA (blue) and ST (orange). Vertical dashed line represents the mean for the corresponding colored distribution.

**Supplementary Figure 3.**

Models trained with overlapping genes and brain areas between ST and ABA datasets are evaluated within dataset on the test fold and across dataset on the entire opposite dataset as illustrated in Figure 1C. (A) Summary diagrams showing mean AUROC for within dataset test set performance (purple arrow) and cross dataset performance with models trained in the opposite dataset (light blue arrow) using the average of 100 feature sets chosen with CFS.

Distributions of AUROCs for within (purple) and cross dataset (light blue) performance for 100 averaged CFS picked gene sets trained (B) in ST and (C) in ABA. In both plots, dashed vertical lines represent the mean of the corresponding colored distribution.

Summary diagram showing mean AUROCs using (D) linear regression and (E) LASSO (alpha = 0.05) for separating out the two planes of slicing in the ABA and treating them alongside the ST dataset as three different datasets for cross-dataset learning. In all three summary diagrams (A, D, E), cross dataset arrows originate from the dataset that the model is trained in and point to the dataset that those models are tested in.

**Supplementary Figure 4.**

Cross-dataset AUROCs (x-axis) of classifying all leaf brain areas from all other leaf brain areas for (A) ABA using LASSO (alpha = 0.05) trained in ST, (B) ABA using linear regression trained in ST, (C) ST using LASSO (alpha = 0.05) trained in ABA, and (D) ST using linear regression trained in ABA as a function of path length (x-axis) in the ARA naming hierarchy between the two brain areas being classified.

The same AUROCs (y-axis) from (A-D) shown in (E-H) respectively as a function of minimum Euclidean distance between the two brain areas in the ARA (x-axis). Euclidean distance on the x-axis is binned into deciles for visualization.

All plots show mean AUROCs (points) with standard deviation (vertical bars).

**Supplementary Figure 5.**

LASSO (alpha = 0.01) one v. all test set performance (x-axis) versus average one v. one performance (y-axis) of the same dataset (A) in ABA and (B) in ST.

(C-D) Same as (A and B), but one v. one performance (y-axis) is accessed in the opposite dataset for (C) train one v. all in ABA and test one v. one in ST and (D) train one v. all in ST and test one v. one in ABA.

(E-H) Same as (A-D) respectively, but using LASSO (alpha = 0.05).

**Supplementary Table 1.**

Examples of brain area pairs from LASSO (alpha=0.1) trained in ST and tested in ABA with minimum path lengths with low and high AUROCs.

**Supplementary Table 2.**

Examples of brain area pairs from LASSO (alpha=0.1) trained in ABA and tested in ST with minimum path lengths with low and high AUROCs.

Supplementary Figure 1

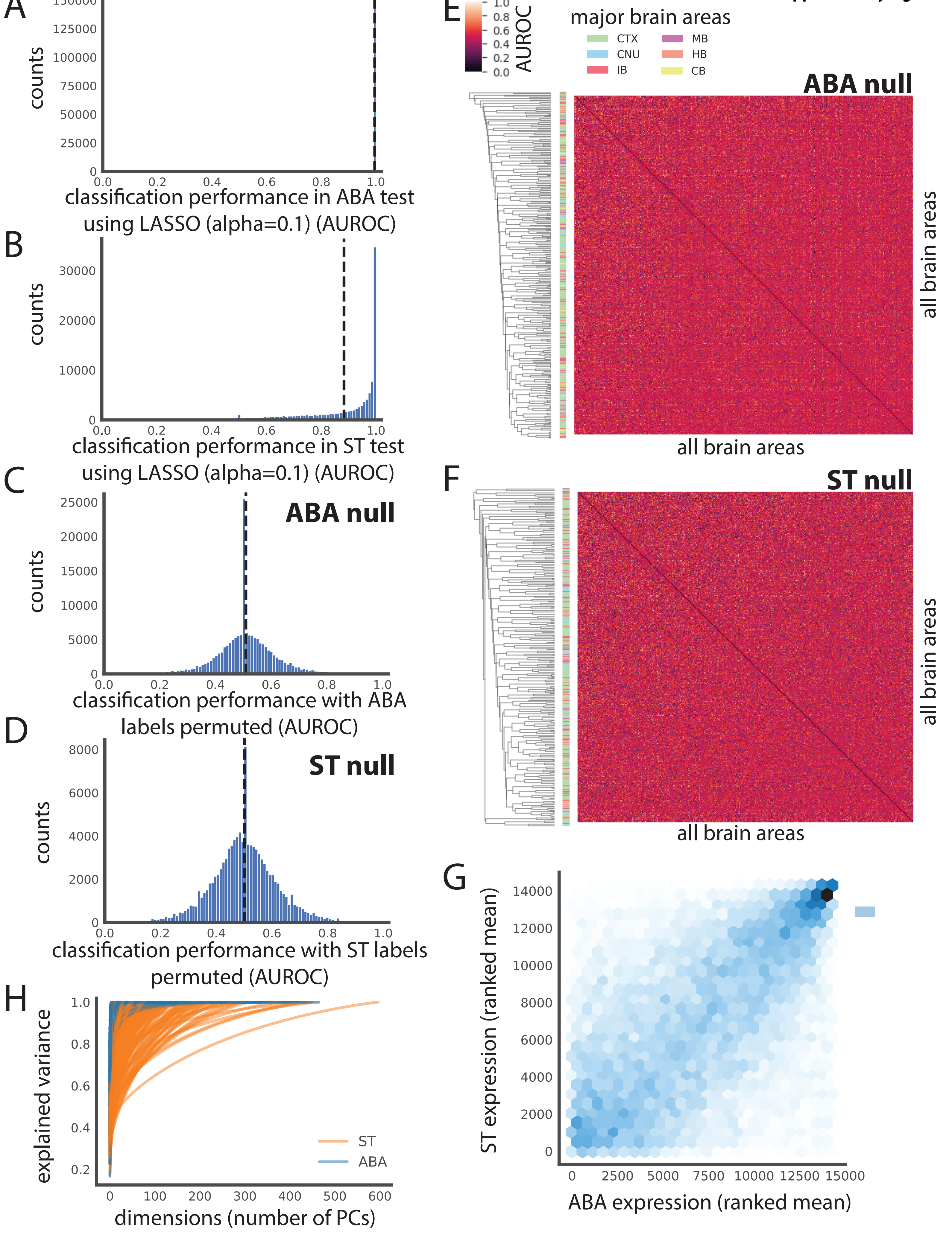

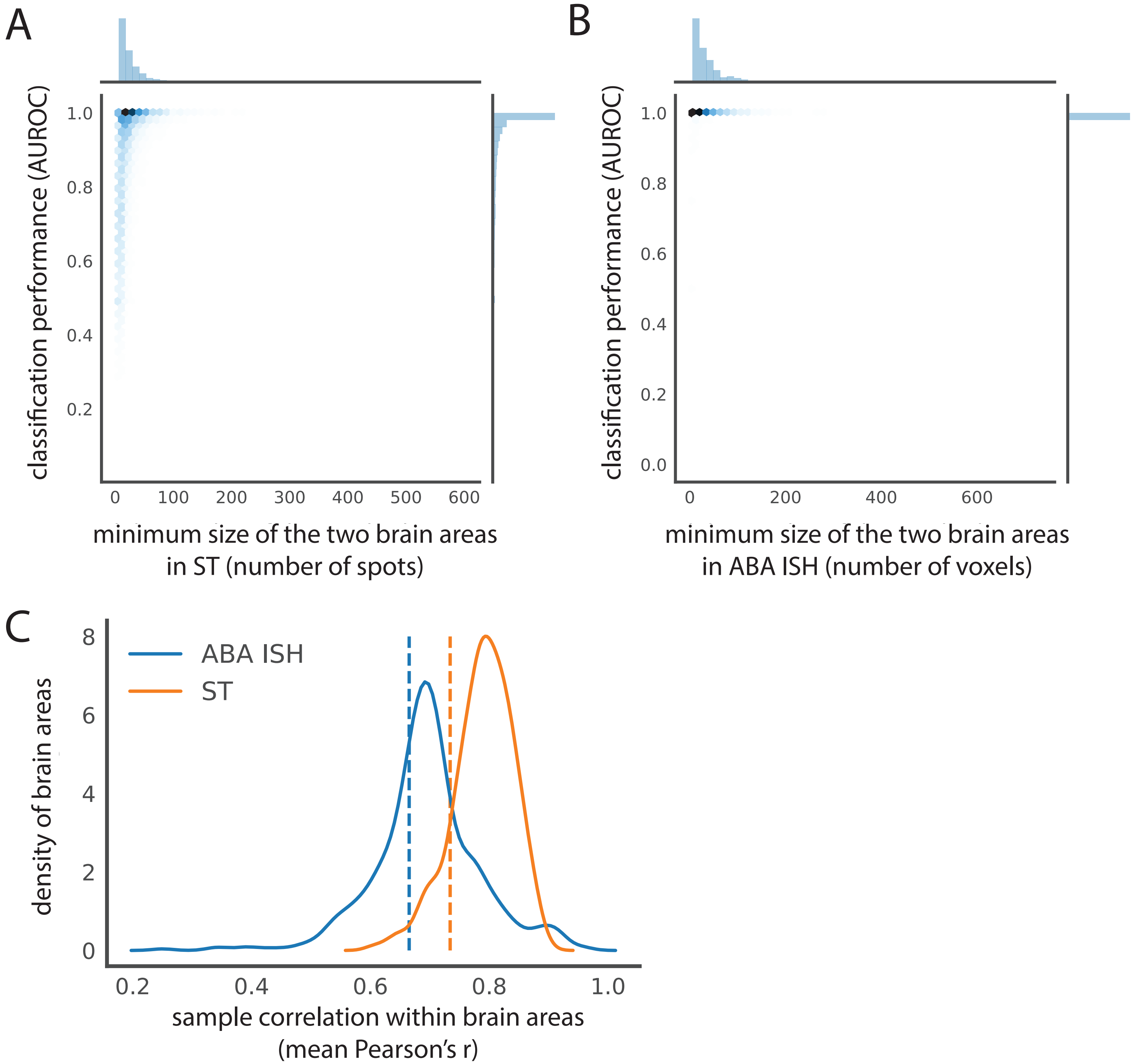


**Supplementary Figure 2**

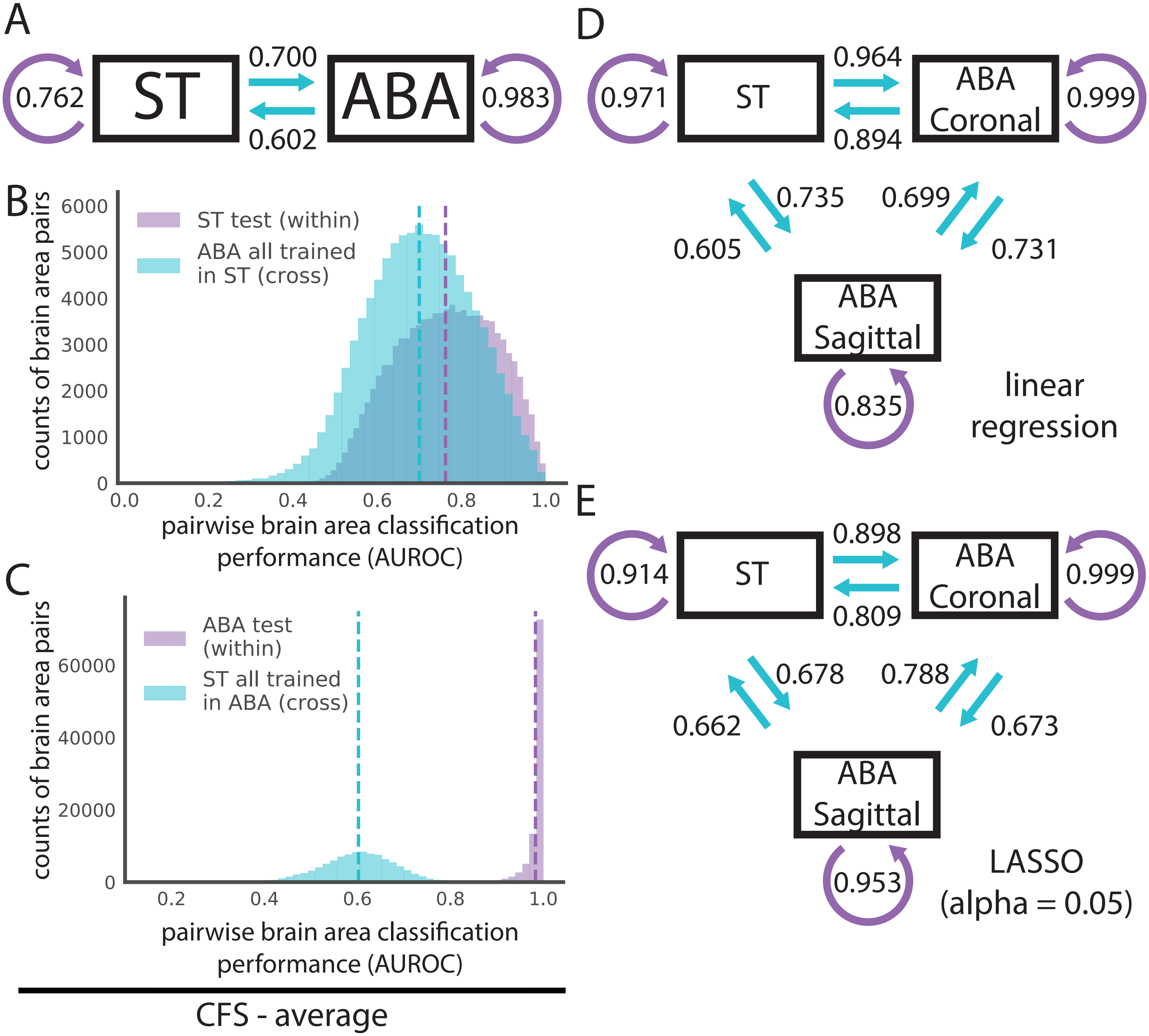


**Supplementary Figure 3**

**Supplementary Figure 4**

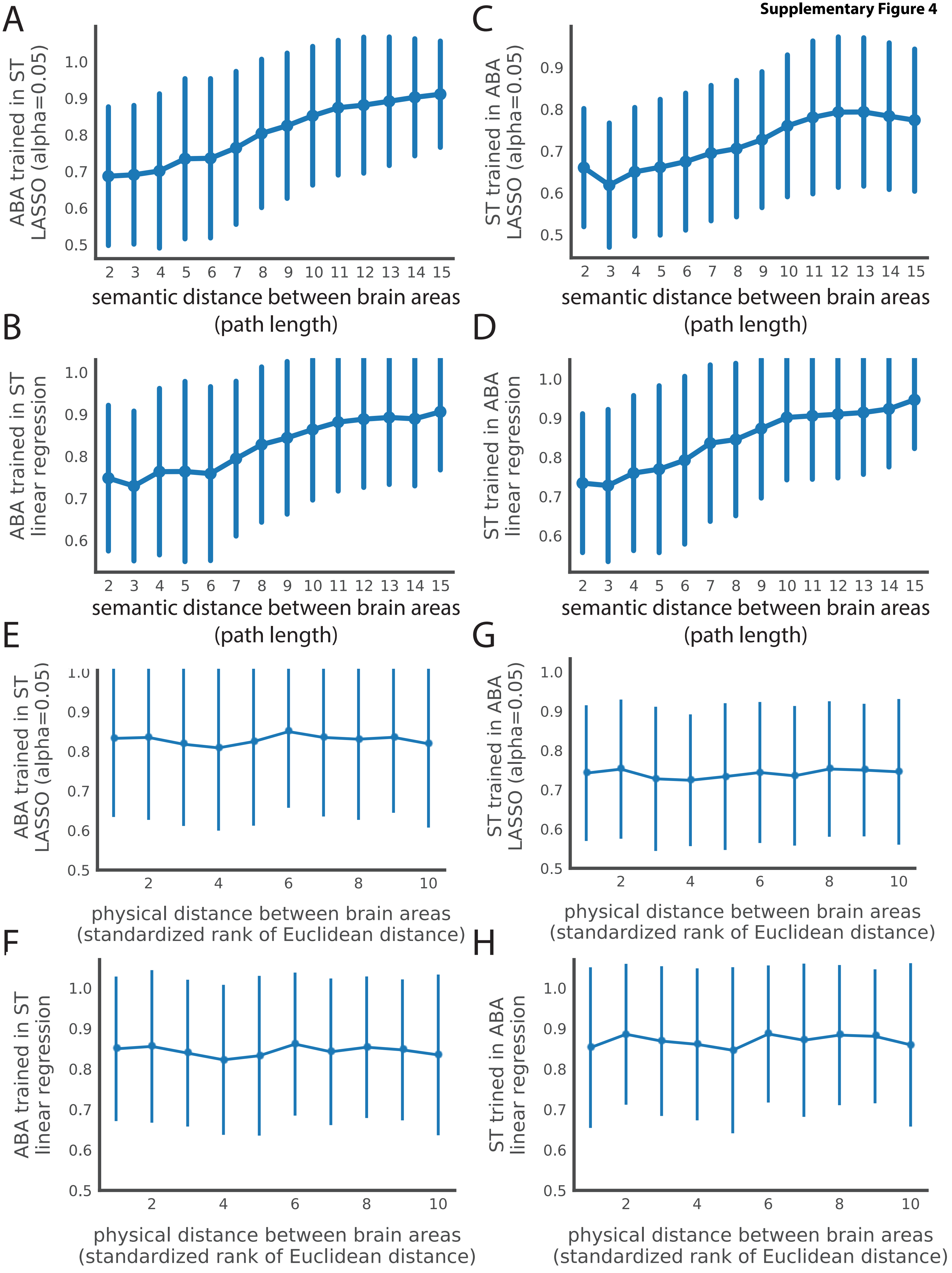

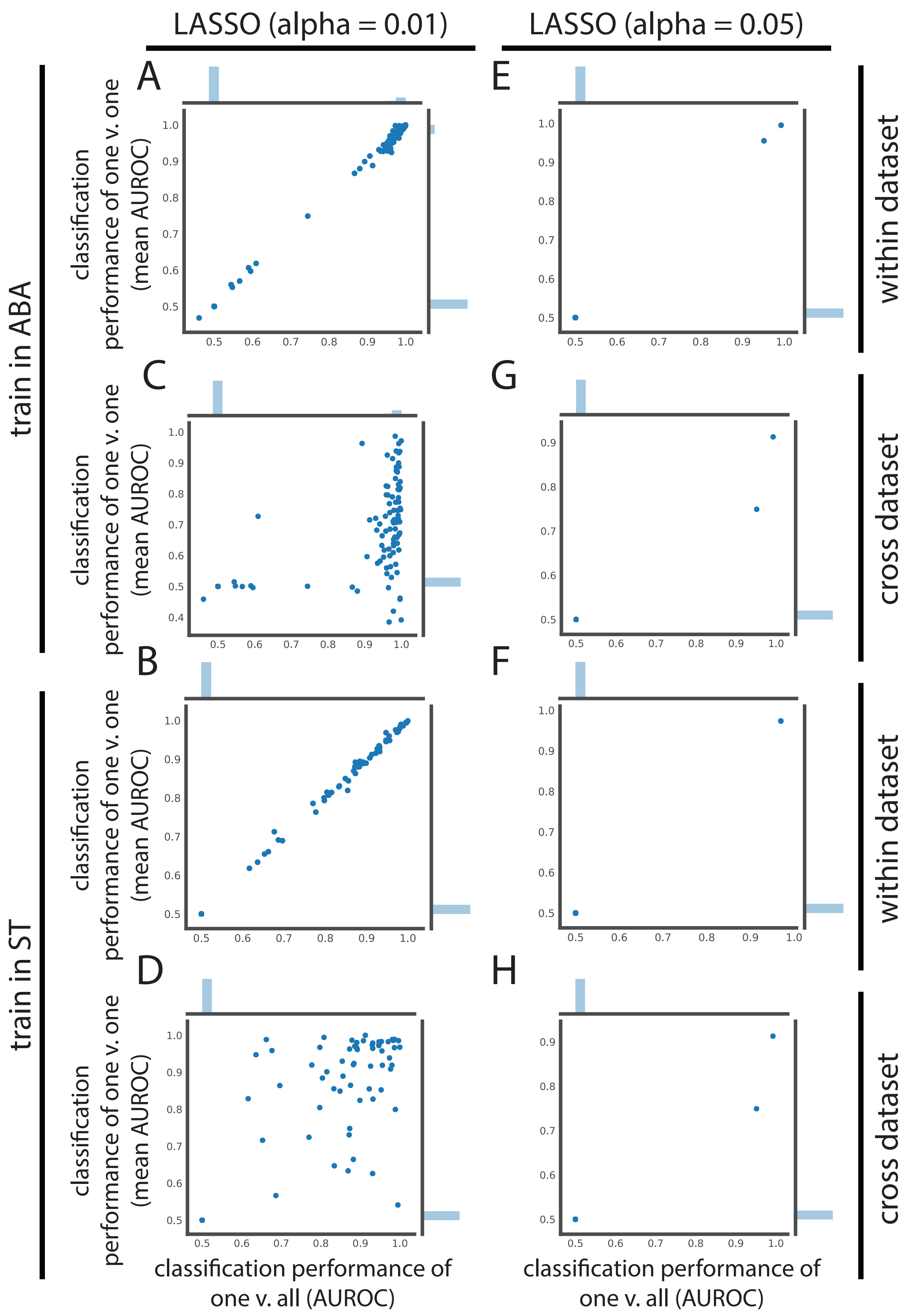


**Supplementary Figure 5**

**Suplementary Table 1**

ST to ABA LASSO, alpha=0.1 auroc file = "STtoABA_ABAall_f1_0p1_051420.csv"

| AUROC = 1; path length = 2 | | | | | | | |
|---|---|---|---|---|---|---|---|
| Brain Area 1 | | | | Brain Area 2 | | | |
| id | acronym | name | parent | id | acronym | name | parent |
| 9 | SSp-tr6a | Primary somatosensory area, trunk, layer 6a | 361 | 1086 | SSp-tr4 | Primary somatosensory area, trunk, layer 4 | 361 |
| 9 | SSp-tr6a | Primary somatosensory area, trunk, layer 6a | 361 | 670 | SSp-tr2/3 | Primary somatosensory area, trunk, layer 2/3 | 361 |
| 1005 | AUDp6b | Primary auditory area, layer 6b | 1002 | 735 | AUDp1 | Primary auditory area, layer 1 | 1002 |
| 1102 | SSp-m6a | Primary somatosensory area, mouth, layer 6a | 345 | 950 | SSp-m4 | Primary somatosensory area, mouth, layer 4 | 345 |
| 1066 | VISam2/3 | Anteromedial visual area, layer 2/3 | 394 | 1046 | VISam6a | Anteromedial visual area, layer 6a | 394 |
| 308 | PTLp6a | Posterior parietal association areas, layer 6a | 22 | 241 | PTLp2/3 | Posterior parietal association areas, layer 2/3 | 22 |
| 1030 | SSp-ll1 | Primary somatosensory area, lower limb, layer 1 | 337 | 478 | SSp-ll6a | Primary somatosensory area, lower limb, layer 6a | 337 |
| 905 | VISal2/3 | Anterolateral visual area, layer 2/3 | 402 | 601 | VISal6a | Anterolateral visual area, layer 6a | 402 |
| 600 | AUDd2/3 | Dorsal auditory area, layer 2/3 | 1011 | 156 | AUDd6a | Dorsal auditory area, layer 6a | 1011 |
| 251 | AUDp2/3 | Primary auditory area, layer 2/3 | 1002 | 847 | AUDp5 | Primary auditory area, layer 5 | 1002 |
| 583 | CLA | Claustrum | 703 | 780 | PA | Posterior amygdalar nucleus | 703 |
| 1035 | SSs4 | Supplemental somatosensory area, layer 4 | 378 | 862 | SSs6a | Supplemental somatosensory area, layer 6a | 378 |
| 729 | TEa6a | Temporal association areas, layer 6a | 541 | 97 | TEa1 | Temporal association areas, layer 1 | 541 |
| 478 | SSp-ll6a | Primary somatosensory area, lower limb, layer 6a | 337 | 113 | SSp-ll2/3 | Primary somatosensory area, lower limb, layer 2/3 | 337 |
| 816 | AUDp4 | Primary auditory area, layer 4 | 1002 | 954 | AUDp6a | Primary auditory area, layer 6a | 1002 |

| AUROC <= 0.5; path length = 2 | | | | | | | |
|---|---|---|---|---|---|---|---|
| Brain Area 1 | | | | Brain Area 2 | | | |
| id | acronym | name | parent | id | acronym | name | parent |
| 1114 | VISal4 | Anterolateral visual area, layer 4 | 402 | 1074 | VISal1 | Anterolateral visual area, layer 1 | 402 |
| 606 | RSPv2 | Retrosplenial area, ventral part, layer 2 | 886 | 622 | RSPv6b | Retrosplenial area, ventral part, layer 6b | 886 |
| 472 | MEApd-a | Medial amygdalar nucleus, posterodorsal part, sublayer a | 426 | 480 | MEApd-b | Medial amygdalar nucleus, posterodorsal part, sublayer b | 426 |
| 1072 | MGd | Medial geniculate complex, dorsal part | 475 | 1088 | MGm | Medial geniculate complex, medial part | 475 |
| 1088 | MGm | Medial geniculate complex, medial part | 475 | 1079 | MGv | Medial geniculate complex, ventral part | 475 |
| 980 | PMd | Dorsal premammillary nucleus | 467 | 1004 | PMv | Ventral premammillary nucleus | 467 |
| 559 | CEAm | Central amygdalar nucleus, medial part | 536 | 544 | CEAc | Central amygdalar nucleus, capsular part | 536 |
| 281 | VISam1 | Anteromedial visual area, layer 1 | 394 | 1066 | VISam2/3 | Anteromedial visual area, layer 2/3 | 394 |
| 1042 | TTd2 | Taenia tecta, dorsal part, layer 2 | 597 | 1050 | TTd3 | Taenia tecta, dorsal part, layer 3 | 597 |
| 148 | GU4 | Gustatory areas, layer 4 | 1057 | 187 | GU5 | Gustatory areas, layer 5 | 1057 |
| 148 | GU4 | Gustatory areas, layer 4 | 1057 | 662 | GU6b | Gustatory areas, layer 6b | 1057 |
| 783 | AId6a | Agranular insular area, dorsal part, layer 6a | 104 | 1101 | AId5 | Agranular insular area, dorsal part, layer 5 | 104 |
| 381 | SNr | Substantia nigra, reticular part | 323 | 616 | CUN | Cuneiform nucleus | 323 |
| 74 | VISl6a | Lateral visual area, layer 6a | 409 | 973 | VISl2/3 | Lateral visual area, layer 2/3 | 409 |
| 74 | VISl6a | Lateral visual area, layer 6a | 409 | 421 | VISl1 | Lateral visual area, layer 1 | 409 |
| 416 | PAA2 | Piriform-amygdalar area, pyramidal layer | 788 | 424 | PAA3 | Piriform-amygdalar area, polymorph layer | 788 |
| 868 | PBld | Parabrachial nucleus, lateral division, dorsal lateral part | 881 | 891 | PBlv | Parabrachial nucleus, lateral division, ventral lateral part | 881 |
| 1106 | VISC2/3 | Visceral area, layer 2/3 | 677 | 897 | VISC1 | Visceral area, layer 1 | 677 |
| 194 | LHA | Lateral hypothalamic area | 290 | 173 | RCH | Retrochiasmatic area | 290 |
| 194 | LHA | Lateral hypothalamic area | 290 | 226 | LPO | Lateral preoptic area | 290 |
| 194 | LHA | Lateral hypothalamic area | 290 | 364 | PSTN | Parasubthalamic nucleus | 290 |
| 368 | PERI6b | Perirhinal area, layer 6b | 922 | 692 | PERI5 | Perirhinal area, layer 5 | 922 |
| 368 | PERI6b | Perirhinal area, layer 6b | 922 | 888 | PERI2/3 | Perirhinal area, layer 2/3 | 922 |
| 1102 | SSp-m6a | Primary somatosensory area, mouth, layer 6a | 345 | 2 | SSp-m6b | Primary somatosensory area, mouth, layer 6b | 345 |
| 969 | ORBvl1 | Orbital area, ventrolateral part, layer 1 | 746 | 608 | ORBvl6a | Orbital area, ventrolateral part, layer 6a | 746 |
| 401 | VISam4 | Anteromedial visual area, layer 4 | 394 | 1046 | VISam6a | Anteromedial visual area, layer 6a | 394 |
| 401 | VISam4 | Anteromedial visual area, layer 4 | 394 | 441 | VISam6b | Anteromedial visual area, layer 6b | 394 |
| 540 | PERI1 | Perirhinal area, layer 1 | 922 | 692 | PERI5 | Perirhinal area, layer 5 | 922 |
| 540 | PERI1 | Perirhinal area, layer 1 | 922 | 888 | PERI2/3 | Perirhinal area, layer 2/3 | 922 |
| 189 | RH | Rhomboid nucleus | 51 | 599 | CM | Central medial nucleus of the thalamus | 51 |
| 880 | DTN | Dorsal tegmental nucleus | 987 | 898 | PCG | Pontine central gray | 987 |
| 1066 | VISam2/3 | Anteromedial visual area, layer 2/3 | 394 | 441 | VISam6b | Anteromedial visual area, layer 6b | 394 |
| 255 | AV | Anteroventral nucleus of thalamus | 239 | 1113 | IAD | Interanterodorsal nucleus of the thalamus | 239 |
| 965 | RSPagl2/3 | Retrosplenial area, lateral agranular part, layer 2/3 | 894 | 774 | RSPagl5 | Retrosplenial area, lateral agranular part, layer 5 | 894 |
| 167 | AONd | Anterior olfactory nucleus, dorsal part | 159 | 160 | AON1 | Anterior olfactory nucleus, layer 1 | 159 |
| 167 | AONd | Anterior olfactory nucleus, dorsal part | 159 | 183 | AONl | Anterior olfactory nucleus, lateral part | 159 |
| 308 | PTLp6a | Posterior parietal association areas, layer 6a | 22 | 340 | PTLp6b | Posterior parietal association areas, layer 6b | 22 |
| 272 | AVPV | Anteroventral periventricular nucleus | 141 | 286 | SCH | Suprachiasmatic nucleus | 141 |
| 272 | AVPV | Anteroventral periventricular nucleus | 141 | 523 | MPO | Medial preoptic area | 141 |
| 1081 | ILA6b | Infralimbic area, layer 6b | 44 | 707 | ILA1 | Infralimbic area, layer 1 | 44 |
| 1081 | ILA6b | Infralimbic area, layer 6b | 44 | 827 | ILA5 | Infralimbic area, layer 5 | 44 |
| 1081 | ILA6b | Infralimbic area, layer 6b | 44 | 556 | ILA2/3 | Infralimbic area, layer 2/3 | 44 |
| 263 | AVP | Anteroventral preoptic nucleus | 141 | 126 | PVp | Periventricular hypothalamic nucleus, posterior part | 141 |
| 1093 | PRNc | Pontine reticular nucleus, caudal part | 987 | 534 | SUT | Supratrigeminal nucleus | 987 |
| 240 | COApm1 | Cortical amygdalar area, posterior part, medial zone, layer 1 | 663 | 248 | COApm2 | Cortical amygdalar area, posterior part, medial zone, layer 2 | 663 |
| 687 | RSPv5 | Retrosplenial area, ventral part, layer 5 | 886 | 622 | RSPv6b | Retrosplenial area, ventral part, layer 6b | 886 |
| 139 | ENTl5 | Entorhinal area, lateral part, layer 5 | 918 | 92 | ENTl4 | Entorhinal area, lateral part, layer 4 | 918 |
| 1030 | SSp-ll1 | Primary somatosensory area, lower limb, layer 1 | 337 | 113 | SSp-ll2/3 | Primary somatosensory area, lower limb, layer 2/3 | 337 |

| | | | | | | | |
|---|---|---|---|---|---|---|---|
| 574 | TRN | Tegmental reticular nucleus | 987 | 534 | SUT | Supratrigeminal nucleus | 987 |
| 837 | SUBd-sr | Subiculum, dorsal part, stratum radiatum | 509 | 845 | SUBd-sp | Subiculum, dorsal part, pyramidal layer | 509 |
| 935 | ACAd1 | Anterior cingulate area, dorsal part, layer 1 | 39 | 211 | ACAd2/3 | Anterior cingulate area, dorsal part, layer 2/3 | 39 |
| 897 | VISC1 | Visceral area, layer 1 | 677 | 857 | VISC6a | Visceral area, layer 6a | 677 |
| 575 | CL | Central lateral nucleus of the thalamus | 51 | 599 | CM | Central medial nucleus of the thalamus | 51 |
| 1074 | VISal1 | Anterolateral visual area, layer 1 | 402 | 905 | VISal2/3 | Anterolateral visual area, layer 2/3 | 402 |
| 1074 | VISal1 | Anterolateral visual area, layer 1 | 402 | 233 | VISal5 | Anterolateral visual area, layer 5 | 402 |
| 544 | CEAc | Central amygdalar nucleus, capsular part | 536 | 551 | CEAl | Central amygdalar nucleus, lateral part | 536 |
| 431 | CA2slm | Field CA2, stratum lacunosum-moleculare | 423 | 454 | CA2sr | Field CA2, stratum radiatum | 423 |
| 303 | BLAa | Basolateral amygdalar nucleus, anterior part | 295 | 451 | BLAv | Basolateral amygdalar nucleus, ventral part | 295 |
| 266 | LSv | Lateral septal nucleus, ventral part | 242 | 258 | LSr | Lateral septal nucleus, rostral (rostroventral) part | 242 |
| 527 | AUDd1 | Dorsal auditory area, layer 1 | 1011 | 600 | AUDd2/3 | Dorsal auditory area, layer 2/3 | 1011 |
| 646 | DP5 | Dorsal peduncular area, layer 5 | 814 | 496 | DP1 | Dorsal peduncular area, layer 1 | 814 |
| 118 | PVi | Periventricular hypothalamic nucleus, intermediate part | 157 | 223 | ARH | Arcuate hypothalamic nucleus | 157 |
| 616 | CUN | Cuneiform nucleus | 323 | 214 | RN | Red nucleus | 323 |
| 772 | ACAv5 | Anterior cingulate area, ventral part, layer 5 | 48 | 810 | ACAv6a | Anterior cingulate area, ventral part, 6a | 48 |
| 269 | VISpl2/3 | Posterolateral visual area, layer 2/3 | 425 | 377 | VISpl6a | Posterolateral visual area, layer 6a | 425 |
| 269 | VISpl2/3 | Posterolateral visual area, layer 2/3 | 425 | 902 | VISpl5 | Posterolateral visual area, layer 5 | 425 |
| 486 | CA3so | Field CA3, stratum oriens | 463 | 471 | CA3slm | Field CA3, stratum lacunosum-moleculare | 463 |
| 1075 | TTv2 | Taenia tecta, ventral part, layer 2 | 605 | 1082 | TTv3 | Taenia tecta, ventral part, layer 3 | 605 |
| 501 | VISpm4 | posteromedial visual area, layer 4 | 533 | 257 | VISpm6a | posteromedial visual area, layer 6a | 533 |
| 92 | ENTl4 | Entorhinal area, lateral part, layer 4 | 918 | 999 | ENTl2/3 | Entorhinal area, lateral part, layer 2/3 | 918 |
| 617 | MDc | Mediodorsal nucleus of the thalamus, central part | 362 | 636 | MDm | Mediodorsal nucleus of the thalamus, medial part | 362 |
| 1045 | ECT6b | Ectorhinal area/Layer 6b | 895 | 977 | ECT6a | Ectorhinal area/Layer 6a | 895 |
| 28 | ENTl6a | Entorhinal area, lateral part, layer 6a | 918 | 60 | ENTl6b | Entorhinal area, lateral part, layer 6b | 918 |
| 243 | AUDd6b | Dorsal auditory area, layer 6b | 1011 | 156 | AUDd6a | Dorsal auditory area, layer 6a | 1011 |
| 52 | ENTl3 | Entorhinal area, lateral part, layer 3 | 918 | 999 | ENTl2/3 | Entorhinal area, lateral part, layer 2/3 | 918 |
| 1142 | TR3 | Postpiriform transition area, layers 3 | 566 | 1141 | TR2 | Postpiriform transition area, layers 2 | 566 |
| 10694 | PAR2 | Parasubiculum, layer 2 | 843 | 10695 | PAR3 | Parasubiculum, layer 3 | 843 |
| 712 | ENTm4 | Entorhinal area, medial part, dorsal zone, layer 4 | 926 | 664 | ENTm3 | Entorhinal area, medial part, dorsal zone, layer 3 | 926 |
| 712 | ENTm4 | Entorhinal area, medial part, dorsal zone, layer 4 | 926 | 727 | ENTm5 | Entorhinal area, medial part, dorsal zone, layer 5 | 926 |
| 565 | VISpm5 | posteromedial visual area, layer 5 | 533 | 257 | VISpm6a | posteromedial visual area, layer 6a | 533 |
| 883 | PBls | Parabrachial nucleus, lateral division, superior lateral part | 881 | 891 | PBlv | Parabrachial nucleus, lateral division, ventral lateral part | 881 |
| 1026 | SSp-ul6b | Primary somatosensory area, upper limb, layer 6b | 369 | 945 | SSp-ul6a | Primary somatosensory area, upper limb, layer 6a | 369 |
| 471 | CA3slm | Field CA3, stratum lacunosum-moleculare | 463 | 479 | CA3slu | Field CA3, stratum lucidum | 463 |
| 471 | CA3slm | Field CA3, stratum lacunosum-moleculare | 463 | 495 | CA3sp | Field CA3, pyramidal layer | 463 |
| 872 | DR | Dorsal nucleus raphe | 165 | 100 | IPN | Interpeduncular nucleus | 165 |
| 460 | MEV | Midbrain trigeminal nucleus | 339 | 580 | NB | Nucleus of the brachium of the inferior colliculus | 339 |
| 162 | LDT | Laterodorsal tegmental nucleus | 1117 | 358 | SLD | Sublaterodorsal nucleus | 1117 |
| 757 | VTN | Ventral tegmental nucleus | 323 | 246 | RR | Midbrain reticular nucleus, retrorubral area | 323 |
| 523 | MPO | Medial preoptic area | 141 | 347 | SBPV | Subparaventricular zone | 141 |
| 523 | MPO | Medial preoptic area | 141 | 126 | PVp | Periventricular hypothalamic nucleus, posterior part | 141 |
| 1010 | VISC4 | Visceral area, layer 4 | 677 | 1058 | VISC5 | Visceral area, layer 5 | 677 |
| 1105 | IA | Intercalated amygdalar nucleus | 278 | 23 | AAA | Anterior amygdalar area | 278 |
| 149 | PVT | Paraventricular nucleus of the thalamus | 571 | 15 | PT | Parataenial nucleus | 571 |
| 604 | NI | Nucleus incertus | 1117 | 238 | RPO | Nucleus raphe pontis | 1117 |
| 564 | MS | Medial septal nucleus | 904 | 596 | NDB | Diagonal band nucleus | 904 |
| 860 | PBlc | Parabrachial nucleus, lateral division, central lateral part | 881 | 875 | PBle | Parabrachial nucleus, lateral division, external lateral part | 881 |
| 860 | PBlc | Parabrachial nucleus, lateral division, central lateral part | 881 | 891 | PBlv | Parabrachial nucleus, lateral division, ventral lateral part | 881 |
| 800 | AIv5 | Agranular insular area, ventral part, layer 5 | 119 | 704 | AIv1 | Agranular insular area, ventral part, layer 1 | 119 |
| 907 | PCN | Paracentral nucleus | 51 | 599 | CM | Central medial nucleus of the thalamus | 51 |
| 335 | PERI6a | Perirhinal area, layer 6a | 922 | 888 | PERI2/3 | Perirhinal area, layer 2/3 | 922 |
| 84 | PL6a | Prelimbic area, layer 6a | 972 | 363 | PL5 | Prelimbic area, layer 5 | 972 |
| 724 | AHNp | Anterior hypothalamic nucleus, posterior part | 88 | 708 | AHNc | Anterior hypothalamic nucleus, central part | 88 |
| 347 | SBPV | Subparaventricular zone | 141 | 126 | PVp | Periventricular hypothalamic nucleus, posterior part | 141 |
| 591 | CLI | Central linear nucleus raphe | 165 | 100 | IPN | Interpeduncular nucleus | 165 |
| 973 | VISl2/3 | Lateral visual area, layer 2/3 | 409 | 421 | VISl1 | Lateral visual area, layer 1 | 409 |
| 810 | ACAv6a | Anterior cingulate area, ventral part, 6a | 48 | 588 | ACAv1 | Anterior cingulate area, ventral part, layer 1 | 48 |
| 700 | AHNa | Anterior hypothalamic nucleus, anterior part | 88 | 708 | AHNc | Anterior hypothalamic nucleus, central part | 88 |
| 232 | COApl3 | Cortical amygdalar area, posterior part, lateral zone, layer 3 | 655 | 224 | COApl2 | Cortical amygdalar area, posterior part, lateral zone, layer 2 | 655 |
| 377 | VISpl6a | Posterolateral visual area, layer 6a | 425 | 902 | VISpl5 | Posterolateral visual area, layer 5 | 425 |
| 1125 | ORBvl5 | Orbital area, ventrolateral part, layer 5 | 746 | 288 | ORBvl2/3 | Orbital area, ventrolateral part, layer 2/3 | 746 |
| 845 | SUBd-sp | Subiculum, dorsal part, pyramidal layer | 509 | 829 | SUBd-m | Subiculum, dorsal part, molecular layer | 509 |
| 1015 | ACAd5 | Anterior cingulate area, dorsal part, layer 5 | 39 | 919 | ACAd6a | Anterior cingulate area, dorsal part, layer 6a | 39 |
| 216 | COApl1 | Cortical amygdalar area, posterior part, lateral zone, layer 1 | 655 | 224 | COApl2 | Cortical amygdalar area, posterior part, lateral zone, layer 2 | 655 |
| 613 | VISl5 | Lateral visual area, layer 5 | 409 | 421 | VISl1 | Lateral visual area, layer 1 | 409 |
| 10693 | PAR1 | Parasubiculum, layer 1 | 843 | 10695 | PAR3 | Parasubiculum, layer 3 | 843 |
| 56 | ACB | Nucleus accumbens | 493 | 998 | FS | Fundus of striatum | 493 |
| 10701 | PRE3 | Presubiculum, layer 3 | 1084 | 10700 | PRE2 | Presubiculum, layer 2 | 1084 |
| 664 | ENTm3 | Entorhinal area, medial part, dorsal zone, layer 3 | 926 | 727 | ENTm5 | Entorhinal area, medial part, dorsal zone, layer 5 | 926 |
| 358 | SLD | Sublaterodorsal nucleus | 1117 | 238 | RPO | Nucleus raphe pontis | 1117 |
| 268 | NLOT2 | Nucleus of the lateral olfactory tract, pyramidal layer | 619 | 1139 | NLOT3 | Nucleus of the lateral olfactory tract, layer 3 | 619 |
| 268 | NLOT2 | Nucleus of the lateral olfactory tract, pyramidal layer | 619 | 260 | NLOT1 | Nucleus of the lateral olfactory tract, molecular layer | 619 |

| | | | | | | | |
|---|---|---|---|---|---|---|---|
| 479 | CA3slu | Field CA3, stratum lucidum | 463 | 495 | CA3sp | Field CA3, pyramidal layer | 463 |
| 511 | SCig-c | Superior colliculus, motor related, intermediate gray layer, sublayer c | 10 | 494 | SCig-a | Superior colliculus, motor related, intermediate gray layer, sublayer a | 10 |
| 310 | SF | Septofimbrial nucleus | 275 | 333 | SH | Septohippocampal nucleus | 275 |
| 484 | ORBm1 | Orbital area, medial part, layer 1 | 731 | 620 | ORBm5 | Orbital area, medial part, layer 5 | 731 |
| 638 | GU6a | Gustatory areas, layer 6a | 1057 | 662 | GU6b | Gustatory areas, layer 6b | 1057 |
| 15 | PT | Parataenial nucleus | 571 | 181 | RE | Nucleus of reunions | 571 |
| 478 | SSp-ll6a | Primary somatosensory area, lower limb, layer 6a | 337 | 510 | SSp-ll6b | Primary somatosensory area, lower limb, layer 6b | 337 |
| 598 | AUDv6b | Ventral auditory area, layer 6b | 1018 | 1023 | AUDv5 | Ventral auditory area, layer 5 | 1018 |
| 137 | CSl | Superior central nucleus raphe, lateral part | 679 | 130 | CSm | Superior central nucleus raphe, medial part | 679 |
| 727 | ENTm5 | Entorhinal area, medial part, dorsal zone, layer 5 | 926 | 743 | ENTm6 | Entorhinal area, medial part, dorsal zone, layer 6 | 926 |
| 450 | SSp-ul1 | Primary somatosensory area, upper limb, layer 1 | 369 | 854 | SSp-ul2/3 | Primary somatosensory area, upper limb, layer 2/3 | 369 |
| 1139 | NLOT3 | Nucleus of the lateral olfactory tract, layer 3 | 619 | 260 | NLOT1 | Nucleus of the lateral olfactory tract, molecular layer | 619 |
| 440 | ORBl6a | Orbital area, lateral part, layer 6a | 723 | 630 | ORBl5 | Orbital area, lateral part, layer 5 | 723 |

**Supplementary Table 2**

ABA to ST LASSO, alpha=0.1 auroc file = "ABAtoST_STall_f1_0p1_051420.csv"

| **AUROC >= 0.95; path length = 2** | | | | | | | |
|---|---|---|---|---|---|---|---|
| **Brain Area 1** | | | | **Brain Area 2** | | | |
| id | acronym | name | parent | id | acronym | name | parent |
| 1005 | AUDp6b | Primary auditory area, layer 6b | 1002 | 816 | AUDp4 | Primary auditory area, layer 4 | 1002 |
| 943 | MOp2/3 | Primary motor area, Layer 2/3 | 985 | 882 | MOp6b | Primary motor area, Layer 6b | 985 |
| 882 | MOp6b | Primary motor area, Layer 6b | 985 | 320 | MOp1 | Primary motor area, Layer 1 | 985 |
| 269 | VISpl2/3 | Posterolateral visual area, layer 2/3 | 425 | 377 | VISpl6a | Posterolateral visual area, layer 6a | 425 |
| 1045 | ECT6b | Ectorhinal area/Layer 6b | 895 | 836 | ECT1 | Ectorhinal area/Layer 1 | 895 |

| **AUROC <=0.5; path length = 2** | | | | | | | |
|---|---|---|---|---|---|---|---|
| **Brain Area 1** | | | | **Brain Area 2** | | | |
| id | acronym | name | parent | id | acronym | name | parent |
| 657 | SSp-m2/3 | Primary somatosensory area, mouth, layer 2/3 | 345 | 950 | SSp-m4 | Primary somatosensory area, mouth, layer 4 | 345 |
| 1114 | VISal4 | Anterolateral visual area, layer 4 | 402 | 233 | VISal5 | Anterolateral visual area, layer 5 | 402 |
| 606 | RSPv2 | Retrosplenial area, ventral part, layer 2 | 886 | 622 | RSPv6b | Retrosplenial area, ventral part, layer 6b | 886 |
| 606 | RSPv2 | Retrosplenial area, ventral part, layer 2 | 886 | 430 | RSPv2/3 | Retrosplenial area, ventral part, layer 2/3 | 886 |
| 472 | MEApd-a | Medial amygdalar nucleus, posterodorsal part, sublayer a | 426 | 487 | MEApd-c | Medial amygdalar nucleus, posterodorsal part, sublayer c | 426 |
| 472 | MEApd-a | Medial amygdalar nucleus, posterodorsal part, sublayer a | 426 | 480 | MEApd-b | Medial amygdalar nucleus, posterodorsal part, sublayer b | 426 |
| 980 | PMd | Dorsal premammillary nucleus | 467 | 946 | PH | Posterior hypothalamic nucleus | 467 |
| 980 | PMd | Dorsal premammillary nucleus | 467 | 1004 | PMv | Ventral premammillary nucleus | 467 |
| 296 | ACAv2/3 | Anterior cingulate area, ventral part, layer 2/3 | 48 | 772 | ACAv5 | Anterior cingulate area, ventral part, layer 5 | 48 |
| 148 | GU4 | Gustatory areas, layer 4 | 1057 | 187 | GU5 | Gustatory areas, layer 5 | 1057 |
| 148 | GU4 | Gustatory areas, layer 4 | 1057 | 662 | GU6b | Gustatory areas, layer 6b | 1057 |
| 783 | AId6a | Agranular insular area, dorsal part, layer 6a | 104 | 1101 | AId5 | Agranular insular area, dorsal part, layer 5 | 104 |
| 381 | SNr | Substantia nigra, reticular part | 323 | 616 | CUN | Cuneiform nucleus | 323 |
| 381 | SNr | Substantia nigra, reticular part | 323 | 757 | VTN | Ventral tegmental nucleus | 323 |
| 191 | AONm | Anterior olfactory nucleus, medial part | 159 | 167 | AONd | Anterior olfactory nucleus, dorsal part | 159 |
| 416 | PAA2 | Piriform-amygdalar area, pyramidal layer | 788 | 424 | PAA3 | Piriform-amygdalar area, polymorph layer | 788 |
| 868 | PBld | Parabrachial nucleus, lateral division, dorsal lateral part | 881 | 860 | PBlc | Parabrachial nucleus, lateral division, central lateral part | 881 |
| 868 | PBld | Parabrachial nucleus, lateral division, dorsal lateral part | 881 | 891 | PBlv | Parabrachial nucleus, lateral division, ventral lateral part | 881 |
| 1106 | VISC2/3 | Visceral area, layer 2/3 | 677 | 1058 | VISC5 | Visceral area, layer 5 | 677 |
| 628 | NOT | Nucleus of the optic tract | 1100 | 634 | NPC | Nucleus of the posterior commissure | 1100 |
| 628 | NOT | Nucleus of the optic tract | 1100 | 215 | APN | Anterior pretectal nucleus | 1100 |
| 105 | SOCm | Superior olivary complex, medial part | 398 | 122 | POR | Superior olivary complex, periolivary region | 398 |
| 194 | LHA | Lateral hypothalamic area | 290 | 364 | PSTN | Parasubthalamic nucleus | 290 |
| 1062 | SSp-bfd6b | Primary somatosensory area, barrel field, layer 6b | 329 | 1070 | SSp-bfd5 | Primary somatosensory area, barrel field, layer 5 | 329 |
| 465 | OT2 | Olfactory tubercle, pyramidal layer | 754 | 473 | OT3 | Olfactory tubercle, polymorph layer | 754 |
| 465 | OT2 | Olfactory tubercle, pyramidal layer | 754 | 481 | isl | Islands of Calleja | 754 |
| 1102 | SSp-m6a | Primary somatosensory area, mouth, layer 6a | 345 | 878 | SSp-m1 | Primary somatosensory area, mouth, layer 1 | 345 |
| 1102 | SSp-m6a | Primary somatosensory area, mouth, layer 6a | 345 | 2 | SSp-m6b | Primary somatosensory area, mouth, layer 6b | 345 |
| 189 | RH | Rhomboid nucleus | 51 | 575 | CL | Central lateral nucleus of the thalamus | 51 |
| 694 | AIv2/3 | Agranular insular area, ventral part, layer 2/3 | 119 | 800 | AIv5 | Agranular insular area, ventral part, layer 5 | 119 |
| 344 | AIp5 | Agranular insular area, posterior part, layer 5 | 111 | 314 | AIp6a | Agranular insular area, posterior part, layer 6a | 111 |
| 965 | RSPagl2/3 | Retrosplenial area, lateral agranular part, layer 2/3 | 894 | 774 | RSPagl5 | Retrosplenial area, lateral agranular part, layer 5 | 894 |
| 272 | AVPV | Anteroventral periventricular nucleus | 141 | 523 | MPO | Medial preoptic area | 141 |
| 272 | AVPV | Anteroventral periventricular nucleus | 141 | 347 | SBPV | Subparaventricular zone | 141 |
| 263 | AVP | Anteroventral preoptic nucleus | 141 | 286 | SCH | Suprachiasmatic nucleus | 141 |
| 263 | AVP | Anteroventral preoptic nucleus | 141 | 523 | MPO | Medial preoptic area | 141 |
| 263 | AVP | Anteroventral preoptic nucleus | 141 | 126 | PVp | Periventricular hypothalamic nucleus, posterior part | 141 |
| 458 | OT1 | Olfactory tubercle, molecular layer | 754 | 481 | isl | Islands of Calleja | 754 |
| 687 | RSPv5 | Retrosplenial area, ventral part, layer 5 | 886 | 430 | RSPv2/3 | Retrosplenial area, ventral part, layer 2/3 | 886 |
| 292 | BA | Bed nucleus of the accessory olfactory tract | 278 | 1105 | IA | Intercalated amygdalar nucleus | 278 |
| 635 | PTLp4 | Posterior parietal association areas, layer 4 | 22 | 241 | PTLp2/3 | Posterior parietal association areas, layer 2/3 | 22 |
| 683 | PTLp5 | Posterior parietal association areas, layer 5 | 22 | 241 | PTLp2/3 | Posterior parietal association areas, layer 2/3 | 22 |
| 622 | RSPv6b | Retrosplenial area, ventral part, layer 6b | 886 | 590 | RSPv6a | Retrosplenial area, ventral part, layer 6a | 886 |
| 622 | RSPv6b | Retrosplenial area, ventral part, layer 6b | 886 | 430 | RSPv2/3 | Retrosplenial area, ventral part, layer 2/3 | 886 |
| 1086 | SSp-tr4 | Primary somatosensory area, trunk, layer 4 | 361 | 461 | SSp-tr6b | Primary somatosensory area, trunk, layer 6b | 361 |
| 305 | VISp6b | Primary visual area, layer 6b | 385 | 33 | VISp6a | Primary visual area, layer 6a | 385 |
| 837 | SUBd-sr | Subiculum, dorsal part, stratum radiatum | 509 | 845 | SUBd-sp | Subiculum, dorsal part, pyramidal layer | 509 |
| 544 | CEAc | Central amygdalar nucleus, capsular part | 536 | 551 | CEAl | Central amygdalar nucleus, lateral part | 536 |
| 411 | MEAad | Medial amygdalar nucleus, anterodorsal part | 403 | 418 | MEAav | Medial amygdalar nucleus, anteroventral part | 403 |
| 614 | TU | Tuberal nucleus | 290 | 173 | RCH | Retrochiasmatic area | 290 |
| 614 | TU | Tuberal nucleus | 290 | 226 | LPO | Lateral preoptic area | 290 |
| 187 | GU5 | Gustatory areas, layer 5 | 1057 | 662 | GU6b | Gustatory areas, layer 6b | 1057 |
| 41 | VISpm2/3 | posteromedial visual area, layer 2/3 | 533 | 565 | VISpm5 | posteromedial visual area, layer 5 | 533 |
| 303 | BLAa | Basolateral amygdalar nucleus, anterior part | 295 | 451 | BLAv | Basolateral amygdalar nucleus, ventral part | 295 |
| 654 | SSp-n4 | Primary somatosensory area, nose, layer 4 | 353 | 838 | SSp-n2/3 | Primary somatosensory area, nose, layer 2/3 | 353 |
| 266 | LSv | Lateral septal nucleus, ventral part | 242 | 258 | LSr | Lateral septal nucleus, rostral (rostroventral) part | 242 |
| 304 | PL2/3 | Prelimbic area, layer 2/3 | 972 | 363 | PL5 | Prelimbic area, layer 5 | 972 |
| 646 | DP5 | Dorsal peduncular area, layer 5 | 814 | 360 | DP2/3 | Dorsal peduncular area, layer 2/3 | 814 |
| 412 | ORBl2/3 | Orbital area, lateral part, layer 2/3 | 723 | 448 | ORBl1 | Orbital area, lateral part, layer 1 | 723 |
| 616 | CUN | Cuneiform nucleus | 323 | 757 | VTN | Ventral tegmental nucleus | 323 |

| | | | | | | | |
|---|---|---|---|---|---|---|---|
| 772 | ACAv5 | Anterior cingulate area, ventral part, layer 5 | 48 | 588 | ACAv1 | Anterior cingulate area, ventral part, layer 1 | 48 |
| 286 | SCH | Suprachiasmatic nucleus | 141 | 347 | SBPV | Subparaventricular zone | 141 |
| 486 | CA3so | Field CA3, stratum oriens | 463 | 471 | CA3slm | Field CA3, stratum lacunosum-moleculare | 463 |
| 1075 | TTv2 | Taenia tecta, ventral part, layer 2 | 605 | 1082 | TTv3 | Taenia tecta, ventral part, layer 3 | 605 |
| 692 | PERI5 | Perirhinal area, layer 5 | 922 | 335 | PERI6a | Perirhinal area, layer 6a | 922 |
| 501 | VISpm4 | posteromedial visual area, layer 4 | 533 | 565 | VISpm5 | posteromedial visual area, layer 5 | 533 |
| 233 | VISal5 | Anterolateral visual area, layer 5 | 402 | 649 | VISal6b | Anterolateral visual area, layer 6b | 402 |
| 233 | VISal5 | Anterolateral visual area, layer 5 | 402 | 601 | VISal6a | Anterolateral visual area, layer 6a | 402 |
| 520 | AUDv6a | Ventral auditory area, layer 6a | 1018 | 598 | AUDv6b | Ventral auditory area, layer 6b | 1018 |
| 1045 | ECT6b | Ectorhinal area/Layer 6b | 895 | 977 | ECT6a | Ectorhinal area/Layer 6a | 895 |
| 52 | ENTl3 | Entorhinal area, lateral part, layer 3 | 918 | 715 | ENTl2a | Entorhinal area, lateral part, layer 2a | 918 |
| 473 | OT3 | Olfactory tubercle, polymorph layer | 754 | 481 | isl | Islands of Calleja | 754 |
| 712 | ENTm4 | Entorhinal area, medial part, dorsal zone, layer 4 | 926 | 664 | ENTm3 | Entorhinal area, medial part, dorsal zone, layer 3 | 926 |
| 883 | PBls | Parabrachial nucleus, lateral division, superior lateral part | 881 | 891 | PBlv | Parabrachial nucleus, lateral division, ventral lateral part | 881 |
| 471 | CA3slm | Field CA3, stratum lacunosum-moleculare | 463 | 495 | CA3sp | Field CA3, pyramidal layer | 463 |
| 670 | SSp-tr2/3 | Primary somatosensory area, trunk, layer 2/3 | 361 | 461 | SSp-tr6b | Primary somatosensory area, trunk, layer 6b | 361 |
| 872 | DR | Dorsal nucleus raphe | 165 | 591 | CLI | Central linear nucleus raphe | 165 |
| 460 | MEV | Midbrain trigeminal nucleus | 339 | 580 | NB | Nucleus of the brachium of the inferior colliculus | 339 |
| 537 | BSTal | Bed nuclei of the stria terminalis, anterior division, anterolateral area | 359 | 498 | BSTam | Bed nuclei of the stria terminalis, anterior division, anteromedial area | 359 |
| 1094 | SSp-ll4 | Primary somatosensory area, lower limb, layer 4 | 337 | 510 | SSp-ll6b | Primary somatosensory area, lower limb, layer 6b | 337 |
| 162 | LDT | Laterodorsal tegmental nucleus | 1117 | 358 | SLD | Sublaterodorsal nucleus | 1117 |
| 162 | LDT | Laterodorsal tegmental nucleus | 1117 | 238 | RPO | Nucleus raphe pontis | 1117 |
| 757 | VTN | Ventral tegmental nucleus | 323 | 749 | VTA | Ventral tegmental area | 323 |
| 757 | VTN | Ventral tegmental nucleus | 323 | 246 | RR | Midbrain reticular nucleus, retrorubral area | 323 |
| 757 | VTN | Ventral tegmental nucleus | 323 | 214 | RN | Red nucleus | 323 |
| 889 | SSp-n6a | Primary somatosensory area, nose, layer 6a | 353 | 702 | SSp-n5 | Primary somatosensory area, nose, layer 5 | 353 |
| 149 | PVT | Paraventricular nucleus of the thalamus | 571 | 15 | PT | Parataenial nucleus | 571 |
| 604 | NI | Nucleus incertus | 1117 | 358 | SLD | Sublaterodorsal nucleus | 1117 |
| 307 | MARN | Magnocellular reticular nucleus | 370 | 661 | VII | Facial motor nucleus | 370 |
| 907 | PCN | Paracentral nucleus | 51 | 599 | CM | Central medial nucleus of the thalamus | 51 |
| 649 | VISal6b | Anterolateral visual area, layer 6b | 402 | 601 | VISal6a | Anterolateral visual area, layer 6a | 402 |
| 724 | AHNp | Anterior hypothalamic nucleus, posterior part | 88 | 708 | AHNc | Anterior hypothalamic nucleus, central part | 88 |
| 591 | CLI | Central linear nucleus raphe | 165 | 100 | IPN | Interpeduncular nucleus | 165 |
| 487 | MEApd-c | Medial amygdalar nucleus, posterodorsal part, sublayer c | 426 | 480 | MEApd-b | Medial amygdalar nucleus, posterodorsal part, sublayer b | 426 |
| 232 | COApl3 | Cortical amygdalar area, posterior part, lateral zone, layer 3 | 655 | 224 | COApl2 | Cortical amygdalar area, posterior part, lateral zone, layer 2 | 655 |
| 377 | VISpl6a | Posterolateral visual area, layer 6a | 425 | 902 | VISpl5 | Posterolateral visual area, layer 5 | 425 |
| 454 | CA2sr | Field CA2, stratum radiatum | 423 | 446 | CA2sp | Field CA2, pyramidal layer | 423 |
| 1113 | IAD | Interanterodorsal nucleus of the thalamus | 239 | 155 | LD | Lateral dorsal nucleus of thalamus | 239 |
| 503 | SCig-b | Superior colliculus, motor related, intermediate gray layer, sublayer b | 10 | 511 | SCig-c | Superior colliculus, motor related, intermediate gray layer, sublayer c | 10 |
| 634 | NPC | Nucleus of the posterior commissure | 1100 | 215 | APN | Anterior pretectal nucleus | 1100 |
| 613 | VISl5 | Lateral visual area, layer 5 | 409 | 421 | VISl1 | Lateral visual area, layer 1 | 409 |
| 56 | ACB | Nucleus accumbens | 493 | 998 | FS | Fundus of striatum | 493 |
| 578 | BSTpr | Bed nuclei of the stria terminalis, posterior division, principal nucleus | 367 | 585 | BSTif | Bed nuclei of the stria terminalis, posterior division, interfascicular nucleus | 367 |
| 676 | DMHp | Dorsomedial nucleus of the hypothalamus, posterior part | 830 | 668 | DMHa | Dorsomedial nucleus of the hypothalamus, anterior part | 830 |
| 360 | DP2/3 | Dorsal peduncular area, layer 2/3 | 814 | 496 | DP1 | Dorsal peduncular area, layer 1 | 814 |
| 479 | CA3slu | Field CA3, stratum lucidum | 463 | 495 | CA3sp | Field CA3, pyramidal layer | 463 |
| 511 | SCig-c | Superior colliculus, motor related, intermediate gray layer, sublayer c | 10 | 494 | SCig-a | Superior colliculus, motor related, intermediate gray layer, sublayer a | 10 |
| 778 | VISp5 | Primary visual area, layer 5 | 385 | 721 | VISp4 | Primary visual area, layer 4 | 385 |
| 310 | SF | Septofimbrial nucleus | 275 | 333 | SH | Septohippocampal nucleus | 275 |
| 638 | GU6a | Gustatory areas, layer 6a | 1057 | 662 | GU6b | Gustatory areas, layer 6b | 1057 |
| 764 | ENTl2b | Entorhinal area, lateral part, layer 2b | 918 | 715 | ENTl2a | Entorhinal area, lateral part, layer 2a | 918 |
| 1046 | VISam6a | Anteromedial visual area, layer 6a | 394 | 441 | VISam6b | Anteromedial visual area, layer 6b | 394 |
| 668 | DMHa | Dorsomedial nucleus of the hypothalamus, anterior part | 830 | 684 | DMHv | Dorsomedial nucleus of the hypothalamus, ventral part | 830 |
| 1127 | TEa2/3 | Temporal association areas, layer 2/3 | 541 | 234 | TEa4 | Temporal association areas, layer 4 | 541 |
| 875 | PBle | Parabrachial nucleus, lateral division, external lateral part | 881 | 891 | PBlv | Parabrachial nucleus, lateral division, ventral lateral part | 881 |
| 1096 | AMd | Anteromedial nucleus, dorsal part | 127 | 1104 | AMv | Anteromedial nucleus, ventral part | 127 |
| 440 | ORBl6a | Orbital area, lateral part, layer 6a | 723 | 630 | ORBl5 | Orbital area, lateral part, layer 5 | 723 |
